# Sensitivity of tree species demography to climate and competition across their range

**DOI:** 10.64898/2026.05.03.722548

**Authors:** Willian Vieira, Andrew MacDonald, Dominique Gravel

## Abstract

Theory predicts that demographic performance should peak at the core of species ranges and decrease toward their limits. Yet, empirical correlations between population growth rate and species distribution remain weak for most tree species. Part of the problem may arise from the difficulty of integrating multiple demographic processes across the complex life cycle of a forest, and from the significant variability among individuals and locations. It remains unclear if the mismatch between performance and distribution arises from modelling limitations or if climate is simply a poor predictor of species performance across distributions. Here, rather than asking whether demographic performance correlates with species distributions, we ask how climate and competition jointly shape population growth rate for 31 tree species across eastern North America. By combining flexible nonlinear hierarchical models for growth, survival, and recruitment with explicit uncertainty propagation, we use Integral Projection Models to address key gaps in previous studies. Perturbation analyses revealed that population growth rate was consistently more sensitive to mean annual temperature than to conspecific or heterospecific competition across all species. We further examined how sensitivities to climate and competition varied across species’ thermal ranges. The dominance of climate over competition increased toward both cold and hot range limits, while sensitivity to competition generally declined from cold to hot limits. Notably, these patterns emerged along the continental thermal gradient shared across species rather than within each species’ individual range, suggesting that range-edge demographic responses may arise as a community-level phenomenon. Across species, the largest source of variability remained the local plot conditions captured by random effects, likely reflecting differences in soil conditions, drainage, and disturbance history. Together, these results may provide a mechanistic pathway underlying the performance declines predicted by range-limit theories, and offer a basis for understanding how forest populations and communities may reorganize in response to ongoing climate change and shifting disturbance regimes.

## 1 Introduction

Mechanistic and process-based models (Evans et al. 2016) have been proposed to improve predictions of species distribution in response to climate change. Among these, demographic range models predict species distributions from individual performance governed by growth, survival, and recruitment rates (Pagel and Schurr 2012). This framework assumes that population growth rate (*λ*), determined by demographic rates, varies across environmental gradients, with species range limits occurring where *λ* is positive (Maguire Jr 1973, Holt 2009, Pang et al. 2024). Approaching species distributions from a demographic perspective allows a more complete representation of how environmental variation and species interactions shape forest dynamics (Schurr et al. 2012, Svenning et al. 2014). Several studies have attempted to predict tree species distributions from demographic performance. The simplest implementations rely on environment-dependent demographic rates to estimate *λ* (e.g. Merow et al. 2014, Csergő et al. 2017). However, competition is a fundamental determinant of demographic rates (Clark et al. 2011, Luo and Chen 2011, Zhang et al. 2015) and population performance (Scherrer et al. 2020, Le Squin et al. 2021) in forest ecosystems. Competition is therefore a necessary addition given the projected risk of community reshuffling under global change (Alexander et al. 2016). This more complete, realized expression of the niche (Hutchinson 1957) may help explain why North American forest trees often fail to occupy their full climatically suitable ranges (Boucher-Lalonde et al. 2012, Talluto et al. 2017).

An increasing body of evidence challenges theoretical expectations by reporting weak correlations between tree demographic performance and species distributions (McGill 2012, Csergő et al. 2017, Bohner and Diez 2020, Le Squin et al. 2021, Midolo et al. 2021, Guyennon et al. 2023, Thuiller et al. 2014). The methodological approaches used to quantify demographic performance are however rarely challenged, difficult to compare and suffer some limitations. Some analyses infer performance from a single proxy, such as radial growth (e.g. McGill 2012, Bohner and Diez 2020). Even when demographic rates are combined within population models, key components such as recruitment are often excluded due to data limitations (Kunstler et al. 2021, Le Squin et al. 2021). Furthermore, density-dependence is sometimes ignored altogether (Csergő et al. 2017, Ohse et al. 2023), and when included, studies rarely distinguish between conspecific and heterospecific competition (Bohner and Diez 2020, Le Squin et al. 2021). Finally, despite increasing recognition of the importance of propagating model and data uncertainty (Milner-Gulland and Shea 2017, Malchow and Hartig 2024), most studies evaluate performance under average environmental conditions and rely on pointwise estimates, missing the substantial variability inherent to demographic processes.

Rather than asking whether demographic performance correlates with species distributions, a more fruitful question may be how climate and competition jointly shape demographic performance. Despite substantial progress, we still lack a full partitioning of the sensitivity of forest dynamics to local and regional drivers (Ohse et al. 2023). For instance, Clark et al. (2011) found that annual growth was more sensitive to competition, whereas fecundity responded more strongly to climate. In contrast, Copenhaver-Parry and Cannon (2016) reported growth to be more sensitive to climate than to competition. Although these studies provide important insights into how forest trees may respond to climate change and management, they focus on individual demographic components rather than their integrated effects on population growth. The distinction matters if sensitivity to climate and competition varies across life-history stages (Russell et al. 2012, Ettinger and HilleRisLambers 2013). The sensitivity of *λ* to climate and competition may also depend on a species’ position within its range. For instance, climate may exert stronger control under abiotic stress, whereas competition may dominate under more benign conditions (Louthan et al. 2015). Yet such range-dependent evaluations of demographic sensitivity remain largely unexplored for forest trees (Ohse et al. 2023).

Here, we evaluate how climate and competition jointly shape the demography and population growth rate of the 31 most abundant forest tree species across eastern North America (Figure 1). Using forest inventories spanning 26–53° latitude across the United States and Canada, we capture the broad geographic ranges of these species. We model growth, survival, and recruitment as functions of temperature, precipitation, and conspecific and heterospecific basal area (as proxies for competition) using Bayesian hierarchical models, with explicit uncertainty propagation. We then integrate these demographic models into size-structured Integral Projection Models (IPMs) to estimate population growth rate (*λ*) under specified climate and competition conditions.

**Figure 1.**
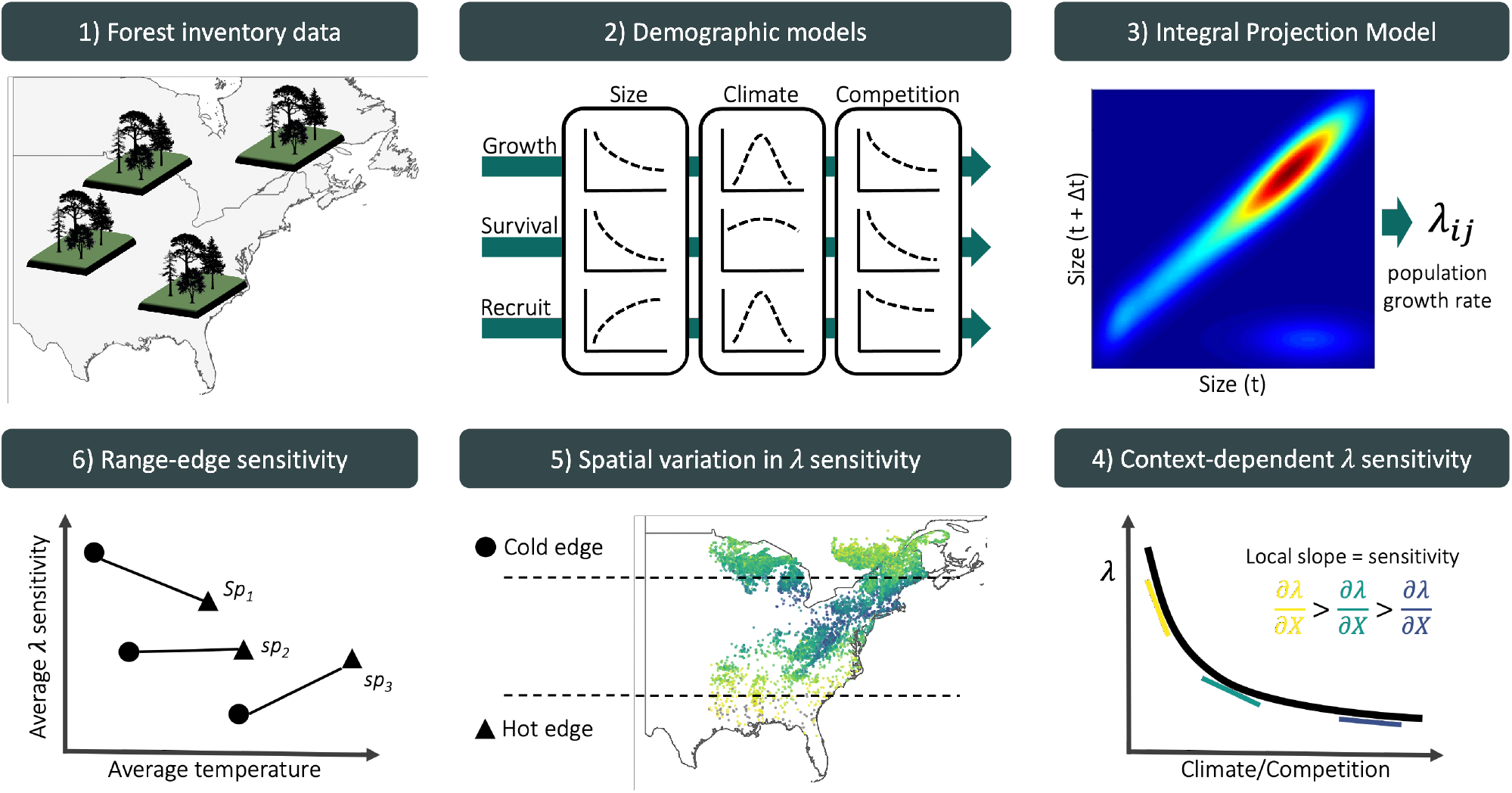
Conceptual framework to assess how the sensitivity of population growth rate to climate and competition varies across species’ geographic ranges. Using repeated measurements of individual trees from forest inventories spanning the eastern United States and Québec, Canada, we fitted species-specific growth, survival, and recruitment models parameterized as functions of individual size, competition, and plot-level climate covariates. These demographic models form the components of a size-structured Integral Projection Model (IPM), from which population growth rate (*λ*) is estimated for each species *i* at each plot location *j*. We then apply perturbation analysis to quantify the sensitivity of *λ* to climate and competition under the local environmental and stand composition conditions of each plot. Cold and hot range limits were identified using the 10th and 90th percentiles of mean annual temperature across each species’ distribution. By averaging sensitivities across plots within each range edge, we assess how the sensitivity of population growth rate differs between cold and hot distributional limits.

Our primary goal is to quantify how sensitive *λ* is to climate and competition across species’ ranges. First, we estimate sensitivities at each plot under observed environmental and competitive conditions using perturbation analysis (Caswell 2000). This approach ensures that sensitivities reflect the environmental conditions actually occupied by the populations. Second, building on previous evidence that North American trees have shown limited cold-edge expansion and hot-edge contraction (Talluto et al. 2017), we test whether demographic sensitivity varies systematically with species’ thermal range position. By quantifying how climate and competition jointly regulate population growth, this framework provides a mechanistic basis for predicting tree responses to climate change, management, and conservation.

## 2 Methods

We fitted species-specific growth, survival, and recruitment models using forest inventory data and integrated them into an Integral Projection Models (IPMs) to estimate population growth rate (*λ*). We then used perturbation analyses to quantify the sensitivity of *λ* to climate and competition across species-specific thermal range positions.

### 2.1 Forest inventory and climate data

We used two open forest inventory datasets from eastern North America: the Forest Inventory and Analysis (FIA) dataset in the United States (O’Connell et al. 2007) and the Forest Inventory of Québec (Ministère des Ressources Naturelles 2016). At the plot level, we focused on plots sampled at least twice and excluded those that had undergone harvesting in order to concentrate solely on natural forest dynamics. Specifically for the FIA dataset, we selected surveys conducted using the modern standardized methodology implemented since 1999. After applying these filters, the final dataset encompassed nearly 26,000 plots spanning a latitudinal range from 26° to 53° (Figure S7). Each plot was measured between 1970 and 2021, with observation frequencies ranging from 2 to 7 measurements and an average of 3 measurements per plot. The time intervals between measurements varied from 1 to 40 years, with a median interval of 7 years (Figure S7).

These datasets provide individual-level information on diameter at breast height (DBH) and survival status (alive or dead) for more than 200 species. From this pool, we selected the 31 most abundant species (Table S1), comprising 9 conifer species and 22 hardwood species. We ensured broad coverage across the shade tolerance gradient, including three very intolerant species, nine intolerant species, eight intermediate species, eight tolerant species, and five very tolerant species (Burns et al. 1990).

We obtained 19 bioclimatic variables at a spatial resolution of 10 km^2^ (300 arcsec), covering the period from 1970 to 2018. These climate variables were modeled using the ANUSPLIN interpolation method (McKenney et al. 2011). Using the geographic coordinates of each plot, we extracted mean annual temperature (MAT) and mean annual precipitation (MAP). When plots did not fall within a valid climate pixel, values were interpolated using the eight neighboring cells. Because measurements occurred across multiple years within census intervals, we calculated both the mean and standard deviation of MAT and MAP across the years included in each time interval.

### 2.2 Model

We developed an Integral Projection Model (IPM) for 31 tree species, with separated sub-models for each demographic rate. An IPM is a mathematical framework used to represent the dynamics of structured populations and communities. Unlike traditional matrix population models, IPMs represent individual traits as continuous variables in discrete time (Easterling et al. 2000). This formulation enables a detailed representation of trait transitions between time steps, which is particularly relevant for trees due to the considerable variability in demographic rates (Kohyama 1992, Clark et al. 2011, Le Squin et al. 2021). Specifically, the IPM predicts the transition of a distribution of individual traits from time *t* to time *t* + Δ*t*, where Δ*t* represents the number of years between observations:

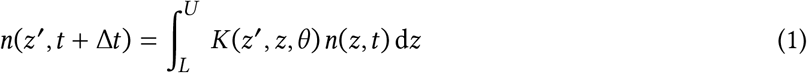

The continuous trait *z* represents DBH, bounded between lower (*L*) and upper (*U*) limits, and *n*(*z, t*) describes the DBH distribution at time *t*. Transitions from *n*(*z, t*) to *n*(*z*^′^, *t* + Δ*t*) are governed by the kernel *K* and species-specific parameters *θ*. The kernel *K*, a continuous analogue of the projection matrix in structured population models, is composed of three demographic components:

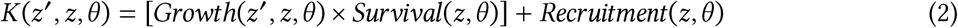

The growth function describes changes in DBH, the survival function determines the probability of remaining alive over the next interval, and the recruitment function describes the influx of new individuals. These three models are estimated independently and do not share parameters. Below, we first describe the baseline versions of these models, followed by the inclusion of climate and competition covariates.

#### 2.2.1 Demographic rates

For the growth model, the DBH of an individual at time *t* + Δ*t* after growing from time *t* follows the distribution:

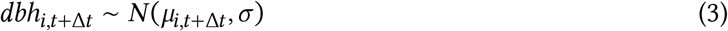

We used the von Bertalanffy growth equation to describe annual DBH growth for individual *i* (Von Bertalanffy 1957). The expected size at time *t* + Δ*t*, given initial size *dbh*_*i,t*_, is:

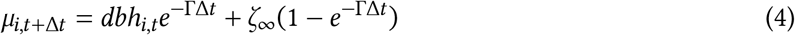

Here, Δ*t* represents the time interval between measurements, Γ is a dimensionless growth-rate parameter, and *ζ*_∞_ represents the asymptotic size at which growth approaches zero. This formulation assumes that growth declines exponentially with size, converging to zero as size approaches *ζ*_∞_. This assumption is particularly valuable in the context of an IPM, as it prevents eviction beyond the bounds of the IPM domain [*L, U*].

For survival, a mortality event (*M*) for individual *i* during the interval between *t* and *t* + Δ*t* is modeled as:

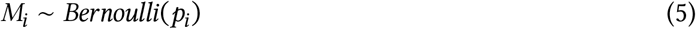

Here, *M*_*i*_ denotes survival status and *p*_*i*_ represents mortality probability. Mortality probability is derived from the annual survival rate (*ψ*):

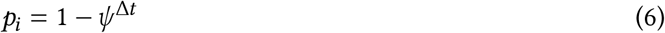

Thus, survival probability increases with longevity parameter *ψ*, while longer intervals Δ*t* increase mortality risk exponentially.

For recruitment, we combined U.S. and Québec inventory data to capture a broader climatic range. However, these inventories differ in protocols for recording seedlings and saplings, including size thresholds. We therefore defined recruitment as the ingrowth of individuals into the adult population (DBH ≥ 12.7 cm).

Recruitment rate (*I*) integrates fecundity, dispersal, growth, and survival up to the threshold size. Because ingrowth depends on census interval length, we defined two parameters: *ϕ*, the annual ingrowth rate per square meter, and *ρ*, the annual survival probability of recruits:

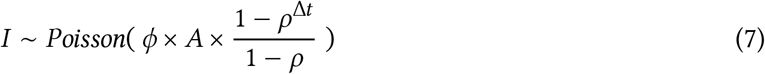

Here, *A* represents plot area (m^2^). Individuals are assumed to enter annually at rate *ϕ*, while their survival (*ρ*) until the subsequent measurement declines with time. Note that *ρ* (recruit survival) is distinct from *ψ* (adult annual survival in Equation 6); the two parameters are estimated independently. *ρ* is estimated from the data of individuals arriving in the population. Once individuals are recruited into the population, their initial size *z*_*I*_ is modeled as:

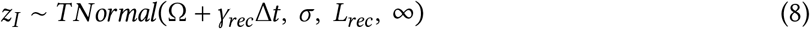

This truncated normal distribution has lower bound *L*_*rec*_ = 12.7 cm and no upper bound. The mean recruit size at observation increases with census interval length at a rate γ_*rec*_.

#### 2.2.2 Covariates

Plot-level random effects were included in all demographic components to account for shared environmental variation among individuals within plots.

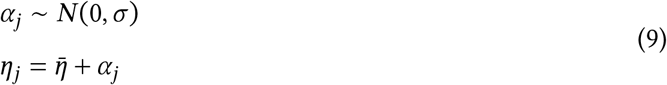

Here, *η* represents parameters Γ, *ψ*, or *ϕ* depending on the demographic component, and *σ* the variance among all plots.

We assumed that competition for light is the primary competitive factor driving forest dynamics (Pacala et al. 1996). We therefore considered that each individual was affected only by larger neighbors. We quantified competition for light for a focal individual in a given plot by summing the basal area of all individuals larger than the focal one, herein BAL. We further split BAL into the total density of conspecific and heterospecific individuals.

Competition effects were modeled on parameters Γ, *ψ*, and *ρ*:

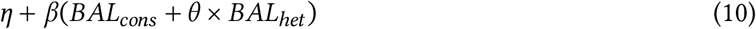

Parameter *θ* controls the relative strength of heterospecific competition. Values *θ* < 1 indicate stronger conspecific competition, *θ* > 1 indicates stronger heterospecific effects, and when *θ* = 1, there is no distinction between them. Note that *β* is also unbounded, allowing it to converge towards negative (indicating competition) or positive (indicating facilitation) values. For recruitment survival (*ρ*), *θ* was fixed at 1 due to convergence issues. Recruitment rate *ϕ* also included conspecific density-dependence, where it decreases with *BAL*_*cons*_ as a positive effect of seed source up to reach the optimal density of recruitment, *δ*, and then decreases with more conspecific density due to competition at a rate proportional to *σ*:

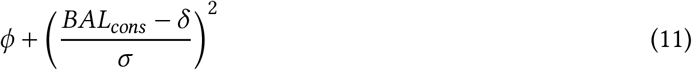

For climate, we used MAT and MAP as climate predictors because of their widespread use in species distribution modeling and demonstrated relevance to model demography of these species (Le Squin et al. 2021). Each demographic parameter followed a unimodal climate response determined by an optimal climate condition (*ξ*) and a climate breadth parameter (*σ*):

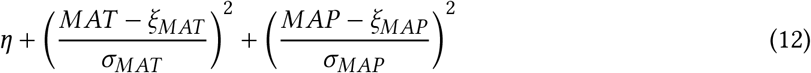

The climate breadth parameter (*σ*) influences the strength of the specific climate variable’s effect on each demographic component. This unimodal function is flexible, assuming various shapes, such as bell, quasi-linear, or flat shapes. However, this flexibility introduces the possibility of parameter degeneracy or redundancy, where different combinations of parameter values yield similar outcomes. To address this issue, we constrained the optimal climate condition parameter (*ξ*) within the observed climate range for the species, assuming that the optimal climate condition falls within our observed data range.

#### 2.2.3 Model fit and validation

We fitted growth, survival, and recruitment models separately for each species using Hamiltonian Monte Carlo (HMC) implemented in Stan (version 2.30.1 Team and Others 2022) through the cmdstanr R interface (version 0.5.3 Gabry et al. 2023). Weakly informative priors were used for all parameters and their full specifications are documented in Table S2. Each model used four chains with 2000 warm-up iterations and 2000 sampling iterations, yielding 8000 posterior samples. To reduce storage, only the final 1000 samples from each chain were retained, resulting in 4000 posterior samples. Models were constructed incrementally, beginning with intercept-only models and progressively adding random effects, competition, and climate covariates. Our goal was not maximal predictive accuracy but mechanistic inference (Tredennick et al. 2021). We focused on assessing the relative effects of climate and competition while controlling for other influential factors. Therefore, our modeling approach is guided by biological mechanisms, which tend to provide more robust extrapolation (Briscoe et al. 2019) rather than solely by statistical metrics. We also verified that increasing model complexity did not reduce predictive performance using mean squared error (MSE), pseudo-*R*^2^ (Gelman et al. 2019), and Leave-One-Out Cross-Validation (LOO-CV). Additional diagnostics are provided in Supplementary Material 1.

The IPM kernel *K* was constructed following Equation 2 and discretized using the midpoint rule (Ellner et al. 2016). Kernel discretization used 0.1 cm size bins, consistent with previous recommendations for trees (Zuidema et al. 2010). The asymptotic population growth rate (*λ*) was computed as the leading eigenvalue of the discretized kernel matrix under specified climate and competition conditions. All model code is available in the `TreesDemography` repository. The IPM implementation is packaged in `forestIPM`, and sensitivity analysis code is available in the `simulations/covariates_perturbation` directory.

### 2.3 Perturbation analysis

Sensitivity was defined as the partial derivative of *λ* with respect to a covariate *X*, representing either competition (conspecific or heterospecific density-dependence) or climate (temperature or precipitation). We quantified sensitivity by slightly increasing each covariate value *X*_*j*_ to 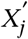 and computing the resulting change in *λ* using a finite-difference approximation (right-hand side of Equation 13):

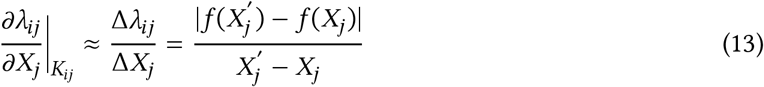

Sensitivity was evaluated separately for each species *i* and plot-year *j*, conditional on the local climate and competition conditions as well as the corresponding kernel parameters *K*_*ij*_. We set the perturbation magnitude to a 1% increase on the normalized scale of each covariate. For instance, a 1% increase corresponds to approximately 0.3°C for mean annual temperature (MAT) and 26 mm for mean annual precipitation (MAP). Because competition metrics were computed at the individual level, perturbations were applied to each individual prior to calculating plot-level basal area, where a 1% increase corresponds approximately to an increase of 1.2 cm in DBH. As we were interested in the magnitude of the response, absolute differences were used, resulting in sensitivity values ranging from 0 to ∞, where lower values indicate weaker sensitivity of *λ* to the given covariate. Specifically, sensitivity (*S*) to competition or climate for species *i* at plot-year *j* was defined as:

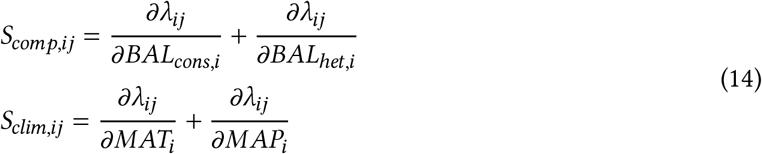

When averaging *S*_*X*,*i*_ across plot-years *j*, this metric represents the sensitivity of *λ*_*i*_ to covariate *X*, conditional on the observed probability distribution of that covariate. This yields a measure of realized sensitivity, integrating both demographic responses and the environmental variability experienced across the species’ range. To evaluate range-dependent sensitivity, plots were categorized into cold, center, or hot conditions along the MAT gradient for each species. Plots were classified as cold (or hot) if their average MAT fell below the 10th percentile (or above the 90th percentile) of the species-specific MAT distribution, with intermediate plots classified as center plots. Sensitivity within each range position was calculated as the average sensitivity across plots belonging to that category. Because this classification is based on observed MAT distributions, range categories are conditional on the environmental conditions experienced by each species.

## 3 Results

### 3.1 Model validation

All species-specific demographic components demonstrated good convergence 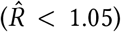 with few to no divergent iterations. Model comparison using LOO-CV consistently favored the full model, featuring plot-level random effects, competition, and climate covariates, over other competing models for all three demographic rates (Supplementary Material 1). The magnitude of LOO-CV differences indicated that the growth model benefited most from the inclusion of covariates, followed by recruitment and survival. We further evaluated biological realism by comparing model-derived parameters with known species trait groups, including growth rate class, maximum observed size, maximum observed age, shade tolerance, and seed mass (Burns et al. 1990, Díaz et al. 2022).

The growth model intercept included two parameters: one determining asymptotic size (*ζ*_∞_) and another describing annual growth rate (Γ). Predicted asymptotic size, which can be interpreted as the maximum predicted size of the species, correlated well with maximum observed species size reported in the literature (*R*^2^ = 0.31, Figure 2). Γ values aligned with established fast-, moderate-, and slow-growing trait classes (Figure S8). In the survival model, expected longevity (*L*) was derived from annual survival probability (*ψ*) using the relationship *L* = *e*^*ψ*^. Estimated longevity showed strong agreement with maximum observed age from the literature (*R*^2^ = 0.59, Figure 2). For recruitment, the log of annual ingrowth rate (*ϕ*) declined linearly with seed mass (Figure S9), consistent with the seed mass–growth rate trade-off (Reich et al. 1998). The annual survival probability of ingrowth (*ρ*) decreased with increasing shade intolerance (Figure S10).

**Figure 2.**
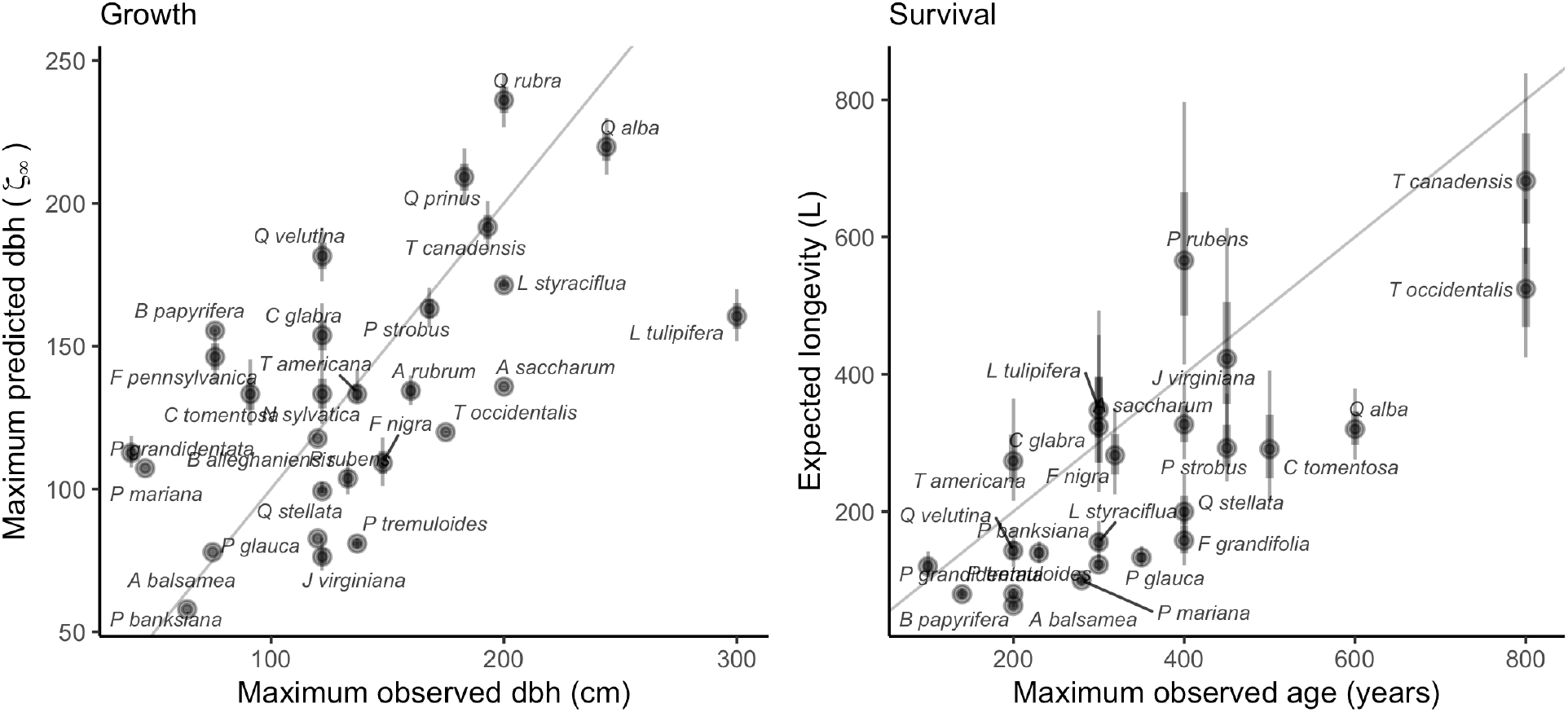
Correlation between predicted asymptotic size (*ζ*_∞_) with maximum observed size (left) and predicted longevity (*L*) with maximum observed age for the 31 forest species. Maximum observed size and age are obtained from Burns et al. (1990). The gray line is the identity curve.

Both conspecific and heterospecific competition effects in growth and survival models increased with shade intolerance (Figure 3). Across nearly all species, conspecific competition effects were stronger than heterospecific effects. Only two species in the growth model and three in the survival model showed stronger heterospecific than conspecific competition. *Fagus grandifolia* and *Thuja occidentalis* were exceptions, exhibiting positive density-dependence in survival. In the recruitment model, total stand density effects increased with shade intolerance (Figure S11).

**Figure 3.**
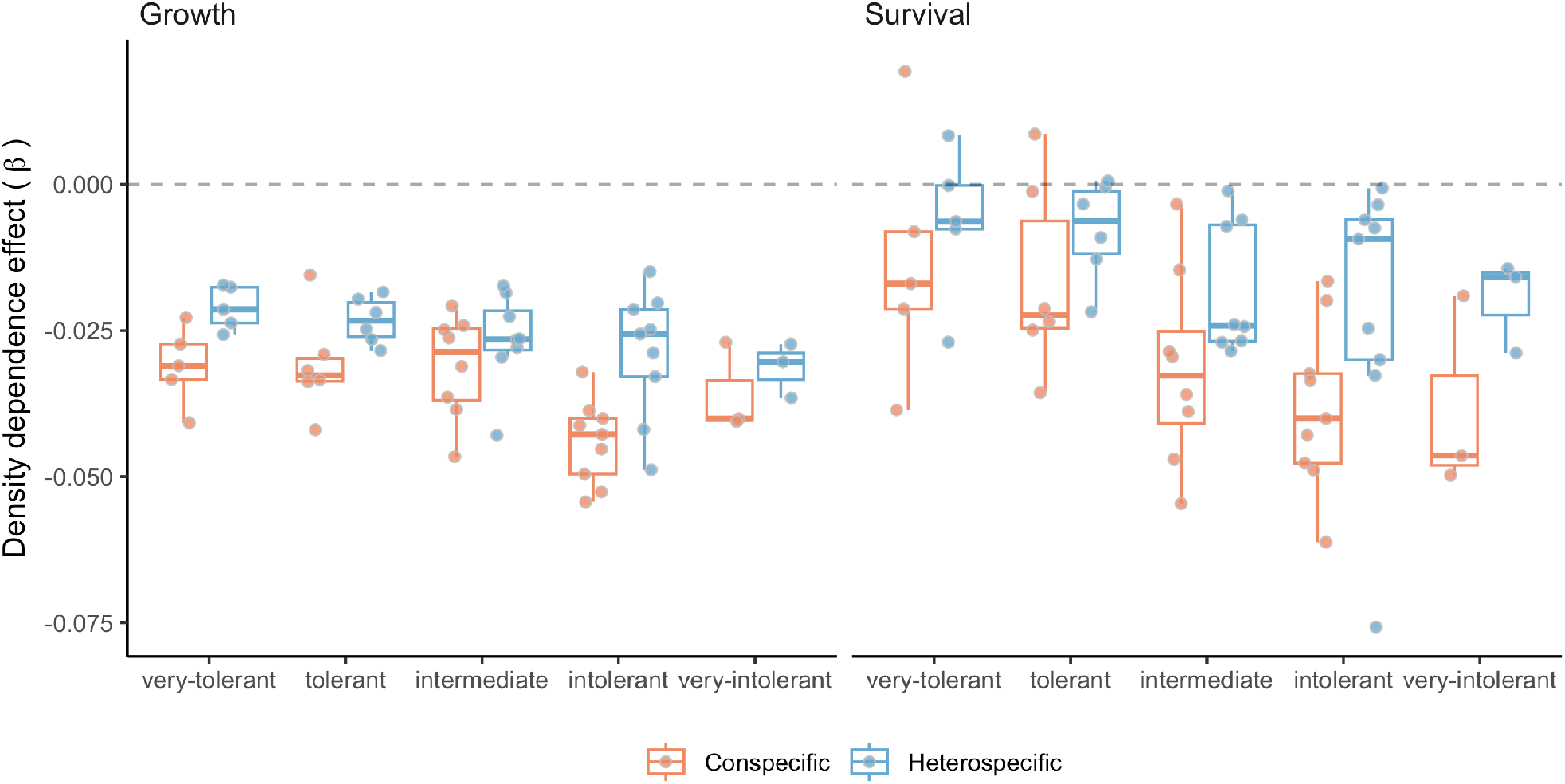
Posterior distribution for the conspecific (red) and heterospecific (blue) density-dependence for each class of shade tolerance (Burns et al. 1990). The more negative the *β*, the stronger the competition effect.

The distribution of optimal MAT (*ξ*_*MAT*_) and MAP (*ξ*_*MAP*_) indicated that optimal conditions for growth, survival, and recruitment were rarely located at the center of species’ ranges (Figures S12 and S13). Many species exhibited demographic compensation, with opposing responses among demographic rates to environmental variation (Villellas et al. 2015). Climate breadth (*σ*) determined how narrowly or broadly performance varied across MAT and MAP. Across species, climate breadth increased with geographic range size, indicating that widely distributed species tended to have broader niche breadths (Figure S14). An exception was survival breadth along MAT, which showed only a weak relationship.

### 3.2 *λ* sensitivity to climate and competition

We conducted perturbation analyses to quantify the relative contribution of each covariate to variation in population growth rate (*λ*). Figure 4 summarizes the sensitivity of *λ* to conspecific competition, heterospecific competition, temperature, and precipitation, averaged across all plot-year observations. Across species, *λ* was most sensitive to temperature, followed by conspecific and heterospecific competition; sensitivity to precipitation was substantially weaker, and this pattern was consistent across species.

**Figure 4.**
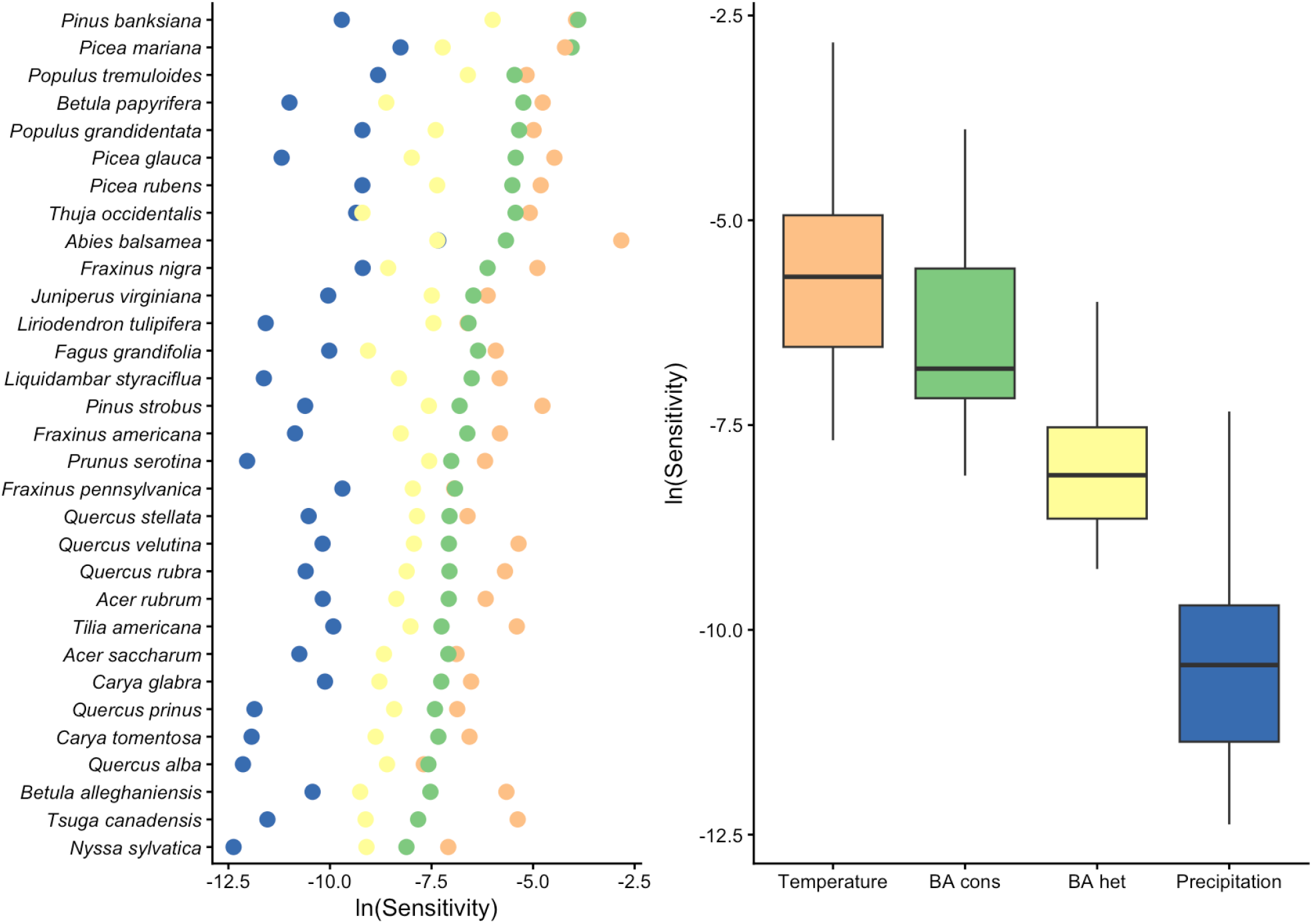
Log sensitivity of species population growth rate to conspecific competition, heterospecific competition, mean annual temperature, and mean annual precipitation across all plot-year observations. The smaller the values, the lower the sensitivity to a covariate.

To evaluate range-dependent responses, plots were divided into cold and hot regions based on species-specific MAT distributions (Figure 5). Most species occurring in colder climates exhibited decreasing climate sensitivity from the cold to the hot limit of their range. In contrast, species centered in warmer climates generally showed greater climate sensitivity at the hot limit than at the cold limit. Sensitivity to competition was typically higher at the cold limit than at the hot limit across most species, regardless of overall distribution. This elevated sensitivity to competition at the cold limit was particularly pronounced among boreal species.

**Figure 5.**
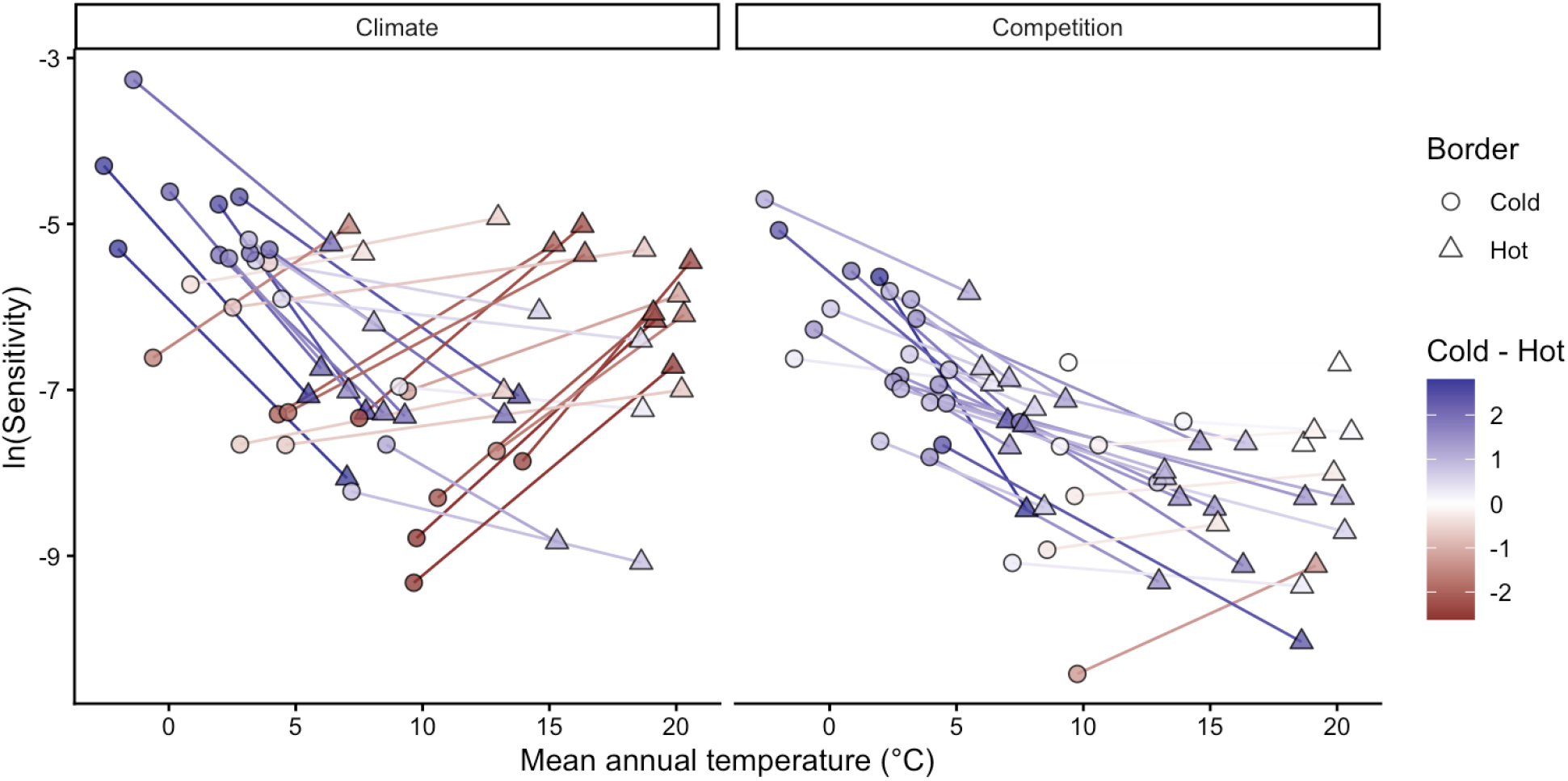
Differences in the sensitivity of species population growth rate to climate (left) and competition between the cold and hot range limits. Each species is represented by a connected line linking their cold (circle) and hot (triangle) range positions, colored according to the difference between the cold and hot sensitivities. Range positions were defined using the median mean annual temperature across all plots belonging to each thermal class. Note that uncertainty in each sensitivity point estimation has been omitted for clarity.

Including sensitivity estimates from the central portion of species ranges further clarified patterns across thermal gradients (Figure S15). Most species displayed approximately linear changes in sensitivity to both climate and competition from cold to hot range limits. For climate sensitivity, three of the four species exhibiting concave relationships, where sensitivity was highest at both range edges, were among those with the largest geographic ranges. In contrast, four species displayed convex relationships in competition sensitivity, with peak sensitivity occurring near the range center. These species also showed the highest overall competition sensitivity and were predominantly distributed in colder climates.

Finally, we evaluated how the relative importance of climate versus competition varied across species’ thermal positions (Figure 6). For most species, *λ* remained more sensitive to climate than to competition across cold, center, and hot range positions. The relative importance of climate increased toward both cold and hot limits along the MAT gradient. Populations near the thermal extremes of the continental climate gradient were therefore more strongly climate-sensitive than those located near the center of the continental climate space. However, the mechanisms underlying this increase differed between range limits (Figure S16). At the cold limit, sensitivities to both climate and competition increased, but the proportional increase was greater for climate. In contrast, at the hot limit, the increasing dominance of climate sensitivity primarily resulted from declining sensitivity to competition.

**Figure 6.**
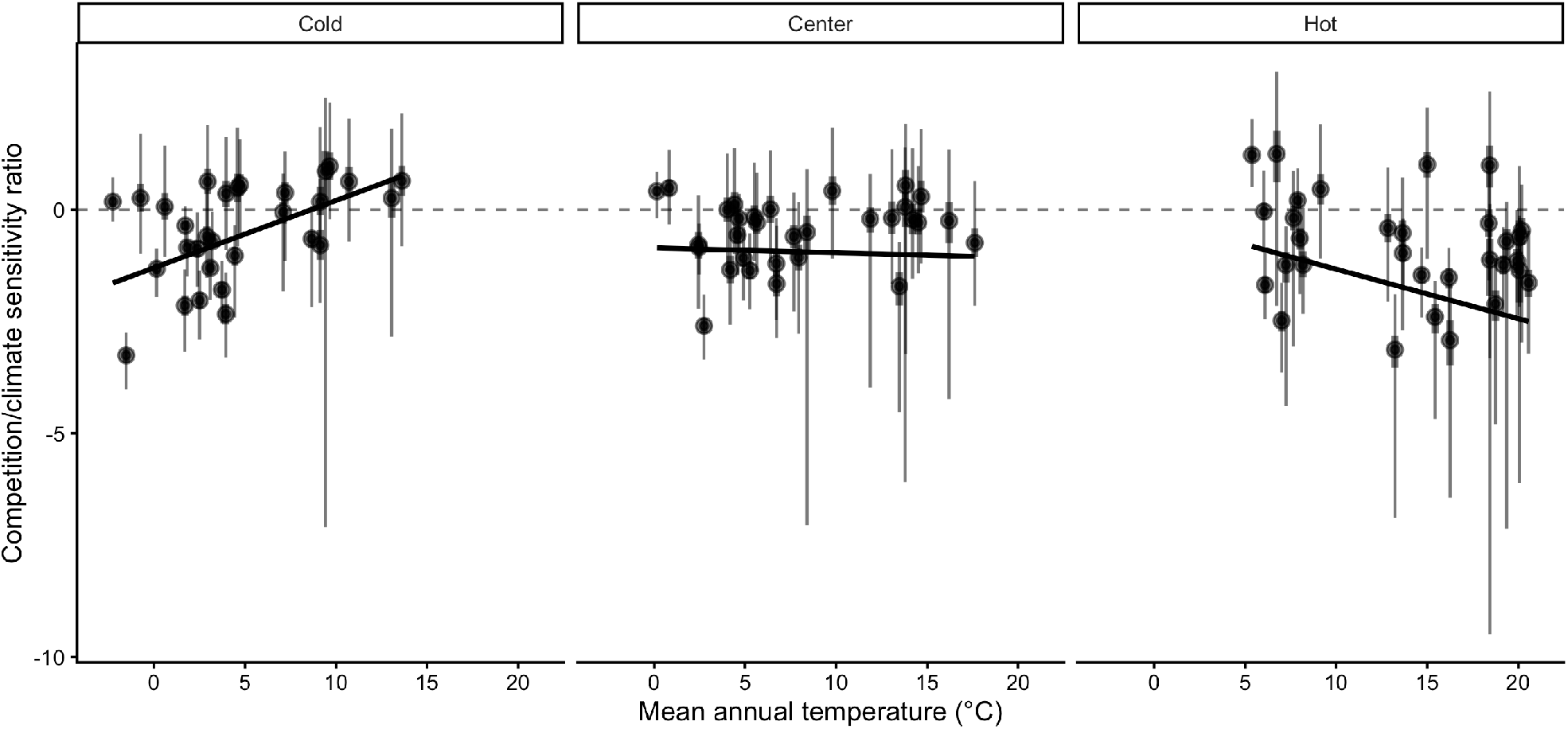
Ratio of population growth rate (*λ*) sensitivity to competition relative to climate across species’ thermal ranges. Negative values indicate greater sensitivity to climate than to competition. Range positions were defined using the median mean annual temperature across all plots belonging to each thermal class. The larger and small bars represent the 20 and 70th quantile probabilities, respectively.

## 4 Discussion

We developed an Integral Projection Model for 31 tree species to evaluate the sensitivity of *λ* to climate and competition. Our framework extends previous analyses of tree species performance by explicitly incorporating climate and competition effects into the recruitment model, distinguishing conspecific from heterospecific competition, and propagating uncertainty across all modeling levels. The modular structure also allows straightforward expansion to the more than 200 species available in the dataset, as well as additional environmental covariates affecting demographic rates.

Adding climate and competition covariates consistently improved predictive performance across all demographic components relative to models containing only random effects. Local plot-level conditions, however, remained the strongest predictors of demographic variation, so we evaluated climate and competition sensitivities while accounting for this plot-level variability. Across species and their geographic ranges, *λ* was most sensitive to temperature and to conspecific basal area of larger individuals. These sensitivities varied across species’ range positions, with climate exerting relatively stronger influence than competition at both cold and hot range limits. Together, these findings contribute to a more mechanistic understanding of how tree species may respond to novel environmental conditions associated with climate change, forest management, and conservation.

### 4.1 Fit of demographic components

Our model demonstrated strong biological consistency by reproducing well-established relationships among demographic traits. Growth and survival intercepts were positively correlated with maximum observed size and longevity, respectively (Burns et al. 1990), while the recruitment intercept showed clear alignment with seed mass (Díaz et al. 2022). The model also reproduced the fast–slow life-history continuum (Salguero-Gómez et al. 2016), reflected in a negative relationship between growth and survival rates and a positive relationship between growth and recruitment rates (Figure S17). Competition effects were consistent with ecological expectations. The model captured the negative relationship between density-dependence and shade tolerance, as well as the general pattern of stronger responses to conspecific competition than to heterospecific competition. This asymmetry is considered fundamental for species coexistence and biodiversity maintenance (Chesson 2000). Conspecific density-dependence was also stronger for fast-growing species than for slow-growing ones (Figure S18), consistent with observations from tropical forests (Zhu et al. 2018). Validation of climate-related parameters remains challenging due to limited empirical estimates of species-specific climatic optima. Nevertheless, our results are aligned with previous findings documenting demographic compensation among forest trees (Bohner and Diez 2020, Yang et al. 2022). The estimated climatic breadth of species also correlated with geographic range size (Figure S14), suggesting that the model captures ecologically meaningful variation beyond explicitly modeled predictors.

Most of the variation in *λ* was associated with local plot conditions captured by random effects, consistent with previous studies (Vanderwel et al. 2016, Le Squin et al. 2021, Itter and Finley 2024). This result indicates that demographic variability is influenced by drivers not explicitly included in our models. At local scales, for instance, soil nitrogen availability (Ibáñez et al. 2018) and mixed mycorrhizal associations (Luo et al. 2023) can enhance tree growth rates. At broader spatial scales, disturbance regimes such as wildfire and insect outbreaks strongly shape forest dynamics and structure (Franklin et al. 2002), causing synchronized mortality events and altering species composition across landscapes. Structured spatial variation in mortality among Pinus species across the United States has similarly been linked primarily to local disturbance regimes rather than broad climatic gradients (Bauman et al. 2025). Although our analysis focused on climate and competition, other covariates may exert stronger influences on demographic variation. For instance, tree growth models incorporating extreme climatic events often show improved predictive performance relative to those using mean climatic conditions (Sanginés de Cárcer et al. 2017). Drought extremes, rather than mean precipitation, have also been identified as strong predictors of fecundity following temperature effects (Clark et al. 2011).

### 4.2 *λ* sensitivity to climate and competition

We found across all species that the sensitivity of *λ* was highest for temperature, followed by conspecific competition. Previous studies assessing the relative importance of climate and competition on tree performance have reported mixed results. Some studies emphasize stronger competition effects on growth (Gómez-Aparicio et al. 2011, Le Squin et al. 2021), whereas others report stronger climate effects (Copenhaver-Parry and Cannon 2016). The relative importance of climate and competition also varies among demographic components, with growth typically more sensitive to competition and fecundity more sensitive to climate (Clark et al. 2011). For instance, several dominant tree species in Poland exhibited high climate sensitivity, with declining fecundity under warming temperatures (Foest et al. 2025). These contrasting findings may arise because many studies assess demographic components separately rather than evaluating their integrated effects on population growth rate. This matters because *λ* does not respond equally to all demographic processes. Our additional analyses revealed that *λ* was most sensitive to recruitment, followed by survival, with a comparatively smaller contribution from growth (Supplementary Material 3). Because recruitment tends to be more sensitive to temperature (Clark et al. 2011, Foest et al. 2025), the high elasticity of *λ* to recruitment may explain the dominant role of climate sensitivity observed in our results.

Demographic sensitivity to climate and competition varied across species’ geographic ranges. Because we modeled climate effects as a unimodal function centered at an optimal value, lower sensitivity values in our framework indicate conditions close to the climatic optimum, whereas higher values indicate deviation from optimal conditions. Overall, sensitivity of *λ* to climate, driven primarily by MAT, was greatest at both cold and hot range limits. This result suggests that species occupying colder portions of the continental thermal gradient have their demographic optimum toward the warmer parts of their ranges, whereas species occupying warmer portions of the continental gradient have their optimum toward the cooler parts. Because multiple species occupy overlapping portions of the same thermal gradient, increasing climatic sensitivity near thermal extremes may represent a shared demographic response to environmental stress rather than an independent property of each species’ geographic range (Körner 2021). This interpretation shifts the focus from species-level range position to community-level responses to climate, highlighting potential common physiological constraints experienced across species.

The demographic mechanisms underlying increased climate sensitivity differed between range limits. Recruitment and growth processes primarily drove sensitivity at cold range limits, whereas survival and recruitment dominated at hot range limits, particularly under high competition conditions (Figure S22). Previous studies have documented climate-constrained growth at cold range limits for both North American (Ettinger and HilleRisLambers 2013) and European trees (Kunstler et al. 2021). Reduced survival at hot range limits has been observed in European forests (Kunstler et al. 2021), although this observation has not been consistently detected in eastern North America (Purves 2009).

Sensitivity of *λ* to competition increased approximately linearly toward colder climates for most species. Due to nonlinear relationships between demographic performance and competition, sensitivity declines as stand density increases, following a negative exponential pattern. The reduction in competition sensitivity toward hot range limits likely reflects higher overall stand density in those regions. Biotic interactions are often thought to dominate at hot range limits (Paquette and Hargreaves 2021). However, when focusing exclusively on growth responses, previous studies have reported relatively constant competition effects across climatic gradients in both North American (Ettinger and HilleRisLambers 2013) and European forests (Kunstler et al. 2011).

### 4.3 Limitations and future perspectives

Structured population models such as IPMs are essential for capturing ontogenetic variation in population dynamics. While our growth model explicitly incorporates individual size, survival and recruitment models were specified without direct size dependence. Attempts to include size-dependent mortality using the widely assumed U-shaped function (Lines et al. 2010) did not improve model performance relative to simpler random-effects formulations (Figure S6). Although mortality often increases with size (Luo and Chen 2011, Hember et al. 2017), its significance appears to manifest only when interacting with climate and competition (Le Squin et al. 2021). The challenge in estimating size-dependent survival likely stems from limited observations of small individuals (dbh < 12.7 cm) and the rarity of large individuals, even in extensive forest inventories (Canham and Murphy 2017). Despite the absence of explicit size dependence in survival, indirect size effects are partially captured through asymmetric competition, where smaller individuals experience stronger competitive pressure. Another limitation shared with many forest-inventory-based models (Kunstler et al. 2021, Le Squin et al. 2021, Guyennon et al. 2023) is the focus on adult trees, even though fecundity can be influenced by climate (Clark et al. 2021), and the dynamics of recruitment may not necessarily align with those of adults (Serra-Diaz et al. 2016, Wason and Dovciak 2017, but see Canham and Murphy 2016).

The modular design of our framework enables direct integration of additional species and environmental predictors. For instance, additional covariates such as water balance or evapotranspiration could be incorporated to assess drought-induced mortality (Peng et al. 2011). Exploring interactions among climate, competition, and individual size may also improve predictions of demographic rates (Peng et al. 2011, Rollinson et al. 2016, Ford et al. 2017, Le Squin et al. 2021). A promising but computationally intensive extension would involve jointly fitting growth, survival, and recruitment models to explicitly capture their interdependence (Pang et al. 2024). Such an approach would allow the incorporation of ecological constraints, such as life-history trade-offs, by sharing information across demographic processes with abundant data (e.g. growth) and those with scarce data (e.g. recruitment). Understanding the ecological drivers underlying variation captured by random effects remains a priority. While our framework accounts for individual and plot-level uncertainty, additional attention to temporal variability in climate and competition will be important. Incorporating temporal stochasticity will improve predictions of species performance under changing environmental conditions and deepen understanding of population responses across space and time (Holt et al. 2022).

## 5 Conclusions

By integrating species-specific growth, survival, and recruitment models into Integral Projection Models, we showed that the population growth rate of 31 eastern North American tree species is consistently more sensitive to mean annual temperature than to competition, with sensitivity to precipitation substantially weaker. Demographic sensitivity varied systematically across thermal range positions: the relative dominance of climate over competition increased toward both cold and hot range limits, with these sensitivity gradients emerging along the continental thermal gradient shared across species rather than within each species’ own range. This result suggests that range-edge demographic responses may emerge as a community-level phenomenon, and that increasing demographic sensitivity toward thermal extremes may provide a mechanistic pathway underlying the expected performance declines predicted by range-limit theories. Most of the variation in *λ*, however, remained tied to local plot conditions captured by random effects, indicating that fine-scale drivers not represented by our climate and competition covariates exert a stronger influence on demography than the broad-scale predictors typically used in range models. Together, these results provide a mechanistic basis for predicting how forest populations may reorganize under climate change, and a framework for understanding the demographic underpinnings of range-limit dynamics.

## Article information

### Open research statement

The data supporting this study were obtained from two open forest inventory programs: the U.S. Forest Inventory and Analysis (FIA) and the Forest Inventory of Québec, Canada. All code to process these datasets and fit the demographic models are available on GitHub at https://github.com/willvieira/TreesDemography. The Integral Projection Model implementation and the perturbation-analysis of this manuscript are available on GitHub at https://github.com/willvieira/forestIPM.

## Acknowledgments

We acknowledge the support of Calcul Québec and Compute Canada for providing the computational resources used in this research. This research was supported by funding from the BIOS^2^ NSERC CREATE program. We are especially grateful to Amaël Le Squin for insightful discussions on the mathematical and Stan models.

## Authorship

Conceptualization: Willian Vieira, Dominique Gravel. Methodology: Willian Vieira, Andrew MacDonald. Formal analysis: Willian Vieira. Visualization: Willian Vieira. Writing — original draft: Willian Vieira. Writing — review and editing: Willian Vieira, Andrew MacDonald, Dominique Gravel.

## Supporting information

Available here.

## References

Alexander, J. M., J. M. Diez, S. P. Hart, and J. M. Levine. 2016. When Climate Reshuffles Competitors: A Call for Experimental Macroecology. Trends in Ecology and Evolution 31:831–841.

Bauman, D., S. M. McMahon, and D. J. Johnson. 2025. Mosaic of size-dependent mortality in three ecologically and economically important pine species reveals patterns across space and time. Global Ecology and Biogeography 34:e70141.

Bohner, T., and J. Diez. 2020. Extensive mismatches between species distributions and performance and their relationship to functional traits. Ecology Letters 23:33–44.

Boucher-Lalonde, V., A. Morin, and D. J. Currie. 2012. How are tree species distributed in climatic space? A simple and general pattern. Global Ecology and Biogeography 21:1157–1166.

Briscoe, N. J., J. Elith, R. Salguero-Gómez, J. J. Lahoz-Monfort, J. S. Camac, K. M. Giljohann, M. H. Holden, B. A. Hradsky, M. R. Kearney, S. M. McMahon, B. L. Phillips, T. J. Regan, J. R. Rhodes, P. A. Vesk, B. A. Wintle, J. D. L. Yen, and G. Guillera-Arroita. 2019. Forecasting species range dynamics with process-explicit models: matching methods to applications. Ecology Letters 22:1940–1956.

Burns, R. M., B. H. Honkala, and Others. 1990. Silvics of North America: 1. Conifers; 2. Hardwoods Agriculture Handbook 654. US Department of Agriculture, Forest Service, Washington, DC.

Canham, C. D., and L. Murphy. 2016. The demography of tree species response to climate: Seedling recruitment and survival. Ecosphere 7:1–16.

Canham, C. D., and L. Murphy. 2017. The demography of tree species response to climate: Sapling and canopy tree survival. Ecosphere 8.

Caswell, H. 2000. Matrix population models. Sinauer Sunderland, MA.

Chesson, P. 2000. Mechanisms of maintenance of species diversity. Annu. Rev. Ecol. Syst 31:343–66.

Clark, J. S., R. Andrus, M. Aubry-Kientz, Y. Bergeron, M. Bogdziewicz, D. C. Bragg, D. Brockway, N. L. Cleavitt, S. Cohen, B. Courbaud, R. Daley, A. J. Das, M. Dietze, T. J. Fahey, I. Fer, J. F. Franklin, C. A. Gehring, G. S. Gilbert, C. H. Greenberg, Q. Guo, J. HilleRisLambers, I. Ibanez, J. Johnstone, C. L. Kilner, J. Knops, W. D. Koenig, G. Kunstler, J. M. LaMontagne, K. L. Legg, J. Luongo, J. A. Lutz, D. Macias, E. J. B. McIntire, Y. Messaoud, C. M. Moore, E. Moran, J. A. Myers, O. B. Myers, C. Nunez, R. Parmenter, S. Pearse, S. Pearson, R. Poulton-Kamakura, E. Ready, M. D. Redmond, C. D. Reid, K. C. Rodman, C. L. Scher, W. H. Schlesinger, A. M. Schwantes, E. Shanahan, S. Sharma, M. A. Steele, N. L. Stephenson, S. Sutton, J. J. Swenson, M. Swift, T. T. Veblen, A. V. Whipple, T. G. Whitham, A. P. Wion, K. Zhu, and R. Zlotin. 2021. Continent-wide tree fecundity driven by indirect climate effects. Nature Communications 12:1–11.

Clark, J. S., D. M. Bell, M. H. Hersh, and L. Nichols. 2011. Climate change vulnerability of forest biodiversity: Climate and competition tracking of demographic rates. Global Change Biology 17:1834–1849.

Copenhaver-Parry, P. E., and E. Cannon. 2016. The relative influences of climate and competition on tree growth along montane ecotones in the Rocky Mountains. Oecologia 182:13–25.

Csergő, A. M., R. Salguero-Gómez, O. Broennimann, S. R. Coutts, A. Guisan, A. L. Angert, E. Welk, I. Stott, B. J. Enquist, B. McGill, J. C. Svenning, C. Violle, and Y. M. Buckley. 2017. Less favourable climates constrain demographic strategies in plants.

Díaz, S., J. Kattge, J. H. C. Cornelissen, I. J. Wright, S. Lavorel, S. Dray, B. Reu, M. Kleyer, C. Wirth, I. C. Prentice, and Others. 2022. The global spectrum of plant form and function: enhanced species-level trait dataset. Scientific Data 9:755.

Easterling, M. R., S. P. Ellner, and P. M. Dixon. 2000. Size-specific sensitivity: applying a new structured population model. Ecology 81:694–708.

Ellner, S. P., D. Z. Childs, and M. Rees. 2016. Data-driven modelling of structured populations. Springer.

Ettinger, A. K., and J. HilleRisLambers. 2013. Climate isn’t everything: Competitive interactions and variation by life stage will also affect range shifts in a warming world. American Journal of Botany 100:1344–1355.

Evans, M. E. K., C. Merow, S. Record, S. M. McMahon, and B. J. Enquist. 2016. Towards Process-based Range Modeling of Many Species. Trends in Ecology and Evolution 31:860–871.

Foest, J. J., J. Szymkowiak, M. Dyderski, D. Kelly, G. Kunstler, S. Jastrzębowski, and M. Bogdziewicz. 2025. Forest fecundity declines as climate shifts.

Ford, K. R., I. K. Breckheimer, J. F. Franklin, J. A. Freund, S. J. Kroiss, A. J. Larson, E. J. Theobald, and J. HilleRisLambers. 2017. Competition alters tree growth responses to climate at individual and stand scales. Canadian Journal of Forest Research 47:53–62.

Franklin, J. F., T. A. Spies, R. V. Pelt, A. B. Carey, D. A. Thornburgh, D. R. Berg, D. B. Lindenmayer, M. E. Harmon, W. S. Keeton, D. C. Shaw, K. Bible, and J. Chen. 2002. Disturbances and structural development of natural forest ecosystems with silvicultural implications, using Douglas-fir forests as an example. Forest Ecology and Management 155:399–423.

Gabry, J., R. Češnovar, and A. Johnson. 2023. cmdstanr: R Interface to ‘CmdStan’.

Gelman, A., B. Goodrich, J. Gabry, and A. Vehtari. 2019. R-squared for Bayesian Regression Models. American Statistician 73:307–309.

Gómez-Aparicio, L., R. García-Valdés, P. Ruíz-Benito, and M. A. Zavala. 2011. Disentangling the relative importance of climate, size and competition on tree growth in Iberian forests: Implications for forest management under global change. Global Change Biology 17:2400–2414.

Guyennon, A., B. Reineking, R. Salguero-Gomez, J. Dahlgren, A. Lehtonen, S. Ratcliffe, P. Ruiz-Benito, M. A. Zavala, and G. Kunstler. 2023. Beyond mean fitness: Demographic stochasticity and resilience matter at tree species climatic edges. Global Ecology and Biogeography 32:573–585.

Hember, R. A., W. A. Kurz, and N. C. Coops. 2017. Relationships between individual-tree mortality and water-balance variables indicate positive trends in water stress-induced tree mortality across North America. Global Change Biology 23:1691–1710.

Holt, R. D. 2009. Bringing the Hutchinsonian niche into the 21st century: Ecological and evolutionary perspectives. Proceedings of the National Academy of Sciences 106:19659–19665.

Holt, R. D., M. Barfield, and J. H. Peniston. 2022. Temporal variation may have diverse impacts on range limits. Philosophical Transactions of the Royal Society B: Biological Sciences 377.

Hutchinson, G. E. 1957. Concluding remarks. Pages 415–427 in Cold spring harbor symposium on quantitative biology.

Ibáñez, I., D. R. Zak, A. J. Burton, and K. S. Pregitzer. 2018. Anthropogenic nitrogen deposition ameliorates the decline in tree growth caused by a drier climate. Ecology.

Itter, M., and A. O. Finley. 2024. Making more with forest inventory data: Toward a scalable, dynamical model of forest change. bioRxiv:2024–07.

Kohyama, T. 1992. Size-structured multi-species model of rain forest trees. Functional Ecology:206–212.

Körner, C. 2021. The cold range limit of trees. Trends in ecology & evolution 36:979–989.

Kunstler, G., C. H. Albert, B. Courbaud, S. Lavergne, W. Thuiller, G. Vieilledent, N. E. Zimmermann, and D. A. Coomes. 2011. Effects of competition on tree radial-growth vary in importance but not in intensity along climatic gradients. Journal of Ecology 99:300–312.

Kunstler, G., A. Guyennon, S. Ratcliffe, N. Rüger, P. Ruiz-Benito, D. Z. Childs, J. Dahlgren, A. Lehtonen, W. Thuiller, C. Wirth, M. A. Zavala, and R. Salguero-Gomez. 2021. Demographic performance of European tree species at their hot and cold climatic edges. Journal of Ecology 109:1041–1054.

Le Squin, A., I. Boulangeat, and D. Gravel. 2021. Climate-induced variation in the demography of 14 tree species is not sufficient to explain their distribution in eastern North America. Global Ecology and Biogeography 30:352–369.

Lines, E. R., D. A. Coomes, and D. W. Purves. 2010. Influences of forest structure, climate and species composition on tree mortality across the Eastern US. PLoS ONE 5.

Louthan, A. M., D. F. Doak, and A. L. Angert. 2015. Where and When do Species Interactions Set Range Limits? Trends in Ecology and Evolution 30:780–792.

Luo, S., R. P. Phillips, I. Jo, S. Fei, J. Liang, B. Schmid, and N. Eisenhauer. 2023. Higher productivity in forests with mixed mycorrhizal strategies. Nature Communications 14:1–10.

Luo, Y., and H. Y. H. Chen. 2011. Competition, species interaction and ageing control tree mortality in boreal forests. Journal of Ecology 99:1470–1480.

Maguire Jr, B. 1973. Niche response structure and the analytical potentials of its relationship to the habitat. The American Naturalist 107:213–246.

Malchow, A.-K., and F. Hartig. 2024. Calibration, sensitivity and uncertainty analysis of ecological models—a review. Vol. in revision). Authorea.

McGill, B. J. 2012. Trees are rarely most abundant where they grow best. Journal of Plant Ecology 5:46–51.

McKenney, D. W., M. F. Hutchinson, P. Papadopol, K. Lawrence, J. Pedlar, K. Campbell, E. Milewska, R. F. Hopkinson, D. Price, and T. Owen. 2011. Customized Spatial Climate Models for North America. Bulletin of the American Meteorological Society 92:1611–1622.

Merow, C., A. M. Latimer, A. M. Wilson, S. M. Mcmahon, A. G. Rebelo, and J. A. Silander. 2014. On using integral projection models to generate demographically driven predictions of species’ distributions: Development and validation using sparse data. Ecography 37:1167–1183.

Midolo, G., C. Wellstein, and S. Faurby. 2021. Individual fitness is decoupled from coarse-scale probability of occurrence in North American trees. Ecography 44:789–801.

Milner-Gulland, E. J., and K. Shea. 2017. Embracing uncertainty in applied ecology. Journal of Applied Ecology.

Ministère des Ressources Naturelles. 2016. Norme d’inventaire ecoforestier: placettes-echantillons temporaires. Direction des inventaires forestier, Ministère des Ressources naturelles,Québec.

O’Connell, M. B., E. B. LaPoint, J. A. Turner, T. Ridley, D. Boyer, A. Wilson, K. L. Waddell, and B. L. Conkling. 2007. The forest inventory and analysis database: Database description and users forest inventory and analysis program. US Department of Agriculture, Forest Service.

Ohse, B., A. Compagnoni, C. E. Farrior, S. M. McMahon, R. Salguero-Gómez, N. Rüger, and T. M. Knight. 2023. Demographic synthesis for global tree species conservation. Trends in Ecology and Evolution 38:579–590.

Pacala, S. W., C. D. Canham, J. Saponara, J. A. Silander, R. K. Kobe, E. Ribbens, J. A. S. Jr., R. K. Kobe, and E. Ribbens. 1996. Forest models defined by Field Measurements: Estimation, Error Analysis and Dynamics. Ecological Monographs 66:1–43.

Pagel, J., and F. M. Schurr. 2012. Forecasting species ranges by statistical estimation of ecological niches and spatial population dynamics. Global Ecology and Biogeography 21:293–304.

Pang, S. E. H., E. Fenollosa, C. Merow, A. Guisan, J.-C. Svenning, and R. Salguero-Gómez. 2024. From niche theory to demographic realities: The demographic niche concept for understanding range-wide population dynamics. Authorea. Under revision in Ecography. doi 10.

Paquette, A., and A. L. Hargreaves. 2021. Biotic interactions are more often important at species’ warm versus cool range edges. Ecology Letters 24:2427–2438.

Peng, C., Z. Ma, X. Lei, Q. Zhu, H. Chen, W. Wang, S. Liu, W. Li, X. Fang, and X. Zhou. 2011. A drought-induced pervasive increase in tree mortality across Canada’s boreal forests. Nature Climate Change 1:467–471.

Purves, D. W. 2009. The demography of range boundaries versus range cores in eastern US tree species. Proceedings of the Royal Society B: Biological Sciences 276:1477–1484.

Reich, P. B., M. G. Tjoelker, M. B. Walters, D. W. Vanderklein, and C. Buschena. 1998. Close association of RGR, leaf and root morphology, seed mass and shade tolerance in seedlings of nine boreal tree species grown in high and low light. Functional Ecology 12:327–338.

Rollinson, C. R., M. W. Kaye, and C. D. Canham. 2016. Interspecific variation in growth responses to climate and competition of five eastern tree species. Ecology 97:1003–1011.

Russell, B. D., C. D. G. Harley, T. Wernberg, N. Mieszkowska, S. Widdicombe, J. M. Hall-Spencer, and S. D. Connell. 2012. Predicting ecosystem shifts requires new approaches that integrate the effects of climate change across entire systems. Biology Letters 8:164–166.

Salguero-Gómez, R., O. R. Jones, E. Jongejans, S. P. Blomberg, D. J. Hodgson, C. Mbeau-Ache, P. A. Zuidema, H. De Kroon, and Y. M. Buckley. 2016. Fast–slow continuum and reproductive strategies structure plant life-history variation worldwide. Proceedings of the National Academy of Sciences 113:230–235.

Sanginés de Cárcer, P., Y. Vitasse, J. Peñuelas, V. E. J. Jassey, A. Buttler, and C. Signarbieux. 2017. Vapor-pressure deficit and extreme climatic variables limit tree growth. Global Change Biology 12:3218–3221.

Scherrer, D., Y. Vitasse, A. Guisan, T. Wohlgemuth, and H. Lischke. 2020. Competition and demography rather than dispersal limitation slow down upward shifts of trees’ upper elevation limits in the Alps. Journal of Ecology:1–15.

Schurr, F. M., J. Pagel, J. S. Cabral, J. Groeneveld, O. Bykova, R. B. O’Hara, F. Hartig, W. D. Kissling, H. P. Linder, G. F. Midgley, B. Schröder, A. Singer, N. E. Zimmermann, R. B. O. Hara, F. Hartig, W. D. Kissling, H. P. Linder, G. F. Midgley, J. W. G.-u. Frankfurt, and F. Main. 2012. How to understand species’ niches and range dynamics: A demographic research agenda for biogeography. Journal of Biogeography 39:2146–2162.

Serra-Diaz, J. M., J. Franklin, W. W. Dillon, A. D. Syphard, F. W. Davis, and R. K. Meentemeyer. 2016. California forests show early indications of both range shifts and local persistence under climate change. Global Ecology and Biogeography 25:164–175.

Svenning, J.-C. C., D. Gravel, R. D. Holt, F. M. Schurr, W. Thuiller, T. Münkemüller, K. H. Schiffers, S. Dullinger, T. C. Edwards, T. Hickler, S. I. Higgins, J. E. M. S. Nabel, J. Pagel, and S. Normand. 2014. The influence of interspecific interactions on species range expansion rates. Ecography 37:1198–1209.

Talluto, M. V., I. Boulangeat, S. Vissault, W. Thuiller, and D. Gravel. 2017. Extinction debt and colonization credit delay range shifts of eastern North American trees. Nature Ecology & Evolution 1:0182.

Team, S. D., and Others. 2022. Stan modeling language users guide and reference manual, version 2.30.1. Stan Development Team.

Thuiller, W., T. Munkemuller, K. H. Schiffers, D. Georges, S. Dullinger, V. M. Eckhart, T. C. Edwards, D. Gravel, G. Kunstler, C. Merow, K. Moore, C. Piedallu, S. Vissault, N. E. Zimmermann, D. Zurell, F. M. Schurr, T. Münkemüller, K. H. Schiffers, D. Georges, S. Dullinger, V. M. Eckhart, T. C. Edwards, D. Gravel, G. Kunstler, C. Merow, K. Moore, C. Piedallu, S. Vissault, N. E. Zimmermann, D. Zurell, and F. M. Schurr. 2014. Does probability of occurrence relate to population dynamics? Ecography 37:1155–1166.

Tredennick, A. T., G. Hooker, S. P. Ellner, and P. B. Adler. 2021. A practical guide to selecting models for exploration, inference, and prediction in ecology. Ecology 102:e03336.

Vanderwel, M. C., H. Zeng, J. P. Caspersen, G. Kunstler, and J. W. Lichstein. 2016. Demographic controls of aboveground forest biomass across North America. Ecology Letters 19:414–423.

Villellas, J., D. F. Doak, M. B. García, and W. F. Morris. 2015. Demographic compensation among populations: What is it, how does it arise and what are its implications? Ecology Letters 18:1139–1152.

Von Bertalanffy, L. 1957. Quantitative laws in metabolism and growth. The quarterly review of biology 32:217–231.

Wason, J. W., and M. Dovciak. 2017. Tree demography suggests multiple directions and drivers for species range shifts in mountains of Northeastern United States. Global Change Biology 23:3335–3347.

Yang, X., A. L. Angert, P. A. Zuidema, F. He, S. Huang, S. Li, S. L. Li, N. I. Chardon, and J. Zhang. 2022. The role of demographic compensation in stabilising marginal tree populations in North America. Ecology Letters 25:1676–1689.

Zhang, J., S. Huang, and F. He. 2015. Half-century evidence from western Canada shows forest dynamics are primarily driven by competition followed by climate. Proceedings of the National Academy of Sciences 112:4009–4014.

Zhu, Y., S. A. Queenborough, R. Condit, S. P. Hubbell, K. P. Ma, and L. S. Comita. 2018. Density-dependent survival varies with species life-history strategy in a tropical forest. Ecology Letters.

Zuidema, P. A., E. Jongejans, P. D. Chien, H. J. During, and F. Schieving. 2010. Integral projection models for trees: a new parameterization method and a validation of model output. Journal of Ecology 98:345– 355.

