## Supporting information for "Sensitivity of tree species demography to climate and competition across their range"

**References** **34**

### 1 Supplementary Material 1

#### 1.1 Model fit

We assessed the growth, survival, and recruitment rates by examining transitions between two measurements. While we fitted the growth and survival functions at the individual level, recruitment was evaluated at the plot level. Due to differences in measurement thresholds between the FIA and Quebec protocols, we only considered individuals with a dbh $\geq$ 127 mm. Therefore, we quantified the ingrowth rate as the number of individuals crossing the 127 mm threshold. We included trees with at least two measurements over time to quantify growth and survival. Similarly, we used plots with at least two measurements over time for ingrowth. To simplify the model hierarchy, we did not incorporate temporal models. Instead, we treated two transition measurements for the same individual as independent information. The plot random effects partially accounted for the variation at the individual level, where different individuals with multiple measurements shared the same variation.

We fitted each of the growth, survival, and recruitment models separately for each species, using the Hamiltonian Monte Carlo (HMC) algorithm via the Stan software (version 2.30.1 Team and Others 2022) and the `cmdstanr` R package (version 0.5.3 Gabry et al. 2023). We conducted 2000 iterations for the warm-up and sampling phases for each of the four chains, resulting in 8000 posterior samples. However, we kept only the last 1000 iterations of the sampling phase to save computation time and storage space, resulting in 4000 posterior samples. We assessed model convergence using Stan's $\hat{R}$ statistic, considering convergence achieved when $\hat{R} < 1.05$. The complete code used for data preparation, model execution, and diagnostic analysis is hosted at https://github.com/willvieira/TreesDemography. Diagnostic reports for all fitted models, including information on model convergence, parameter distributions, prediction checks, $R^2$, and other metrics, are available at https://willvieira.github.io/TreesDemography/.

#### 1.2 Model comparison

We constructed the demographic models incrementally, starting from the simple intercept model and gradually adding plot random effects, competition, and climate covariates. While the intercept-only model represents the most basic form, we opted to discard it and use the intercept model with random effects as the baseline or null model for comparison with more complex models. We ensured the convergence of all these model forms, and comprehensive diagnostic details are available at https://github.com/willvieira/TreesDemography.

Our primary objective is to select the model that has learned the most from the data. We used complementary metrics to quantify the gain in information achieved by adding complexity to each demographic model. One intuitive metric involves assessing the reduction in variance attributed to likelihood and the variance associated with plot random effects. A greater reduction in their variance implies a greater information gain from model complexity. The following metrics are all derived from the idea of increasing predictive accuracy. Although we focus on inference, measuring predictive power is crucial for quantifying the additional information gained from including new covariates. The first two classic measures of predictive accuracy are the mean squared error (MSE) and the pseudo $R^2$. We base these metrics on the linear relationship between observed and predicted demographic outputs. Finally, we used Leave-One-Out Cross-Validation (LOO-CV), which uses the sampled data to estimate the model's out-of-sample predictive accuracy (Vehtari et al. 2017). LOO-CV allows us to assess how well each model describes the observed data and compare competing models to determine which has learned the most from the data.

#### 1.3 Parameter variance

This section describes how the variance attributed to plot random effects changes with increasing model complexity. As we introduce covariates, it is expected that part of the variance in demographic rates, initially attributed to random effects, shifts towards the covariate fixed effects. Therefore, the larger the reduction in variance associated with plot random effects, the more significant the role of covariates in explaining demographic rates. The Figure 1 shows the $\sigma_{plot}$ change with increased model complexity for growth, survival, and recruitment vital rates.

#### 1.4 Model predictive accuracy

We used pseudo $R^2$ and MSE metrics derived from comparing observed and predicted values to evaluate the predictive accuracy of growth and recruitment demographic rates. Higher $R^2$ values and lower MSE indicate better overall model accuracy. The Figures 2 and 3 compare the growth and recruitment models using $R^2$ and MSE, respectively.

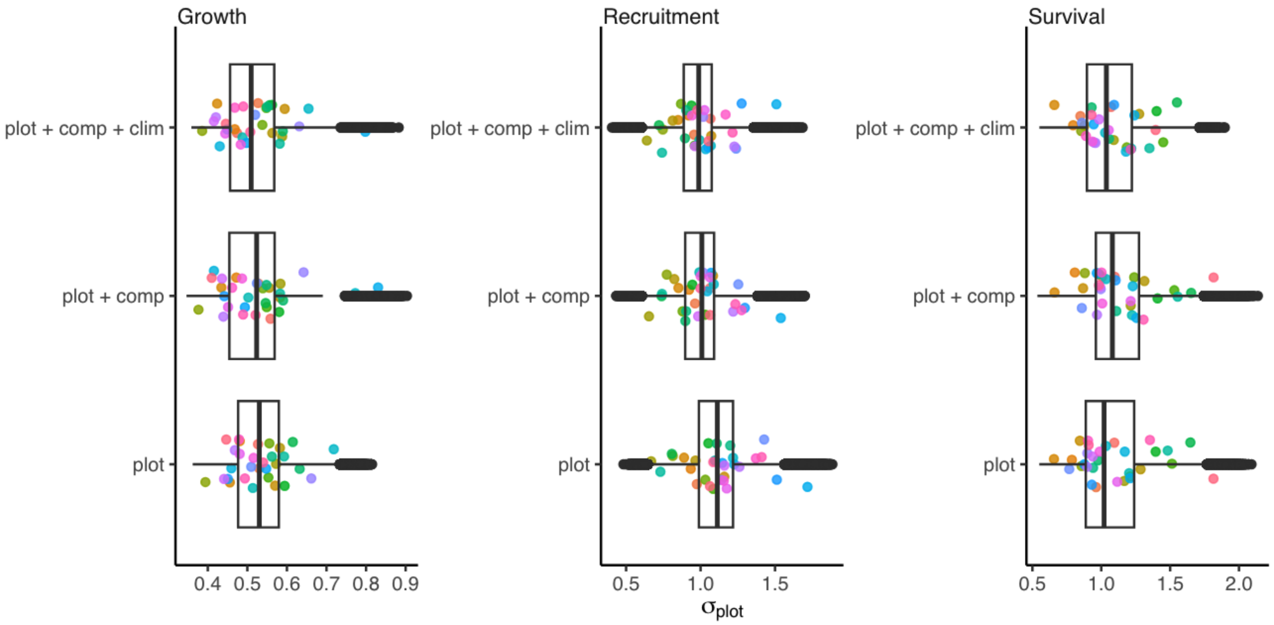

Figure 1: Boxplot showing the change in the posterior distribution of the parameter $\sigma_{plot}$ across the 31 tree species between the competing models. For each growth, survival, and recruitment vital rate, the simplest model (plot random effects only) increases in complexity with the addition of fixed size, competition, and climate covariates. Each colored dot represents the species' average posterior distribution.

We used three complementary metrics for the survival model to assess model predictions. While the accuracy of classification models is often evaluated through the fraction of correct predictions, this measure can be misleading for unbalanced datasets such as mortality, where dead events are rare. To address this issue, we calculated sensitivity, which measures the percentage of dead trees correctly identified as dead (true positives). We also computed specificity, which measures the percentage of live trees correctly identified as alive (true negatives). The combination of sensitivity and specificity allows us to calculate corrected accuracy, considering the unbalanced accuracy predictions of positive and negative events (Figure 4).

#### 1.5 Leave-one-out cross-validation

Finally, we evaluated the competing models using the LOO-CV metric (Figure 5), where models are compared based on the difference in the expected log pointwise predictive density (ELPD_diff). In cases involving multiple models, the difference is calculated relative to the model with highest ELPD (Vehtari et al. 2017). Consequently, the model with ELPD_diff equal to zero is defined as the best model. In contrast, the performance of the other models is assessed based on their deviation from

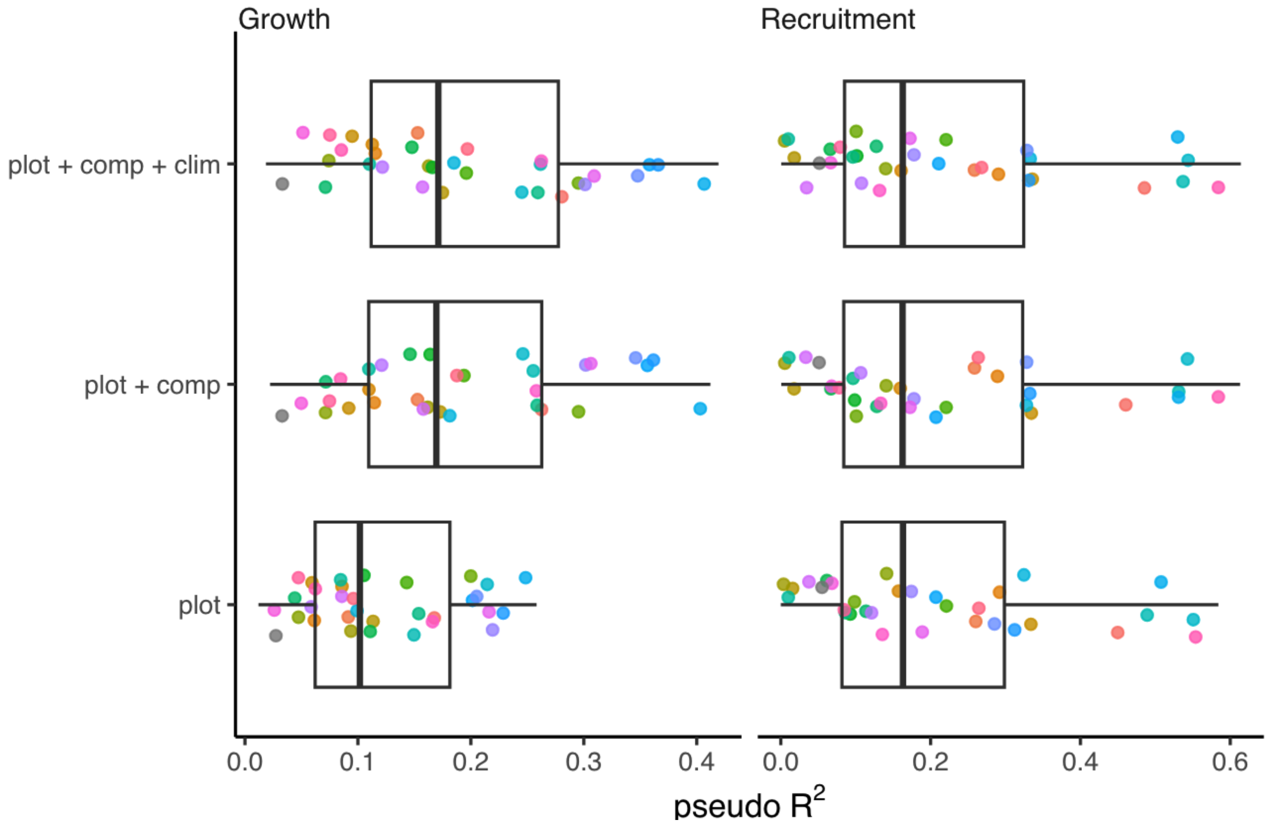

Figure 2: Posterior distribution of pseudo $R^2$ across the 31 tree species between the competing models. For each growth, survival, and recruitment vital rate, the simplest model (plot random effects only) increases in complexity with the addition of fixed competition and climate covariates. Each colored dot represents the species' average posterior distribution.

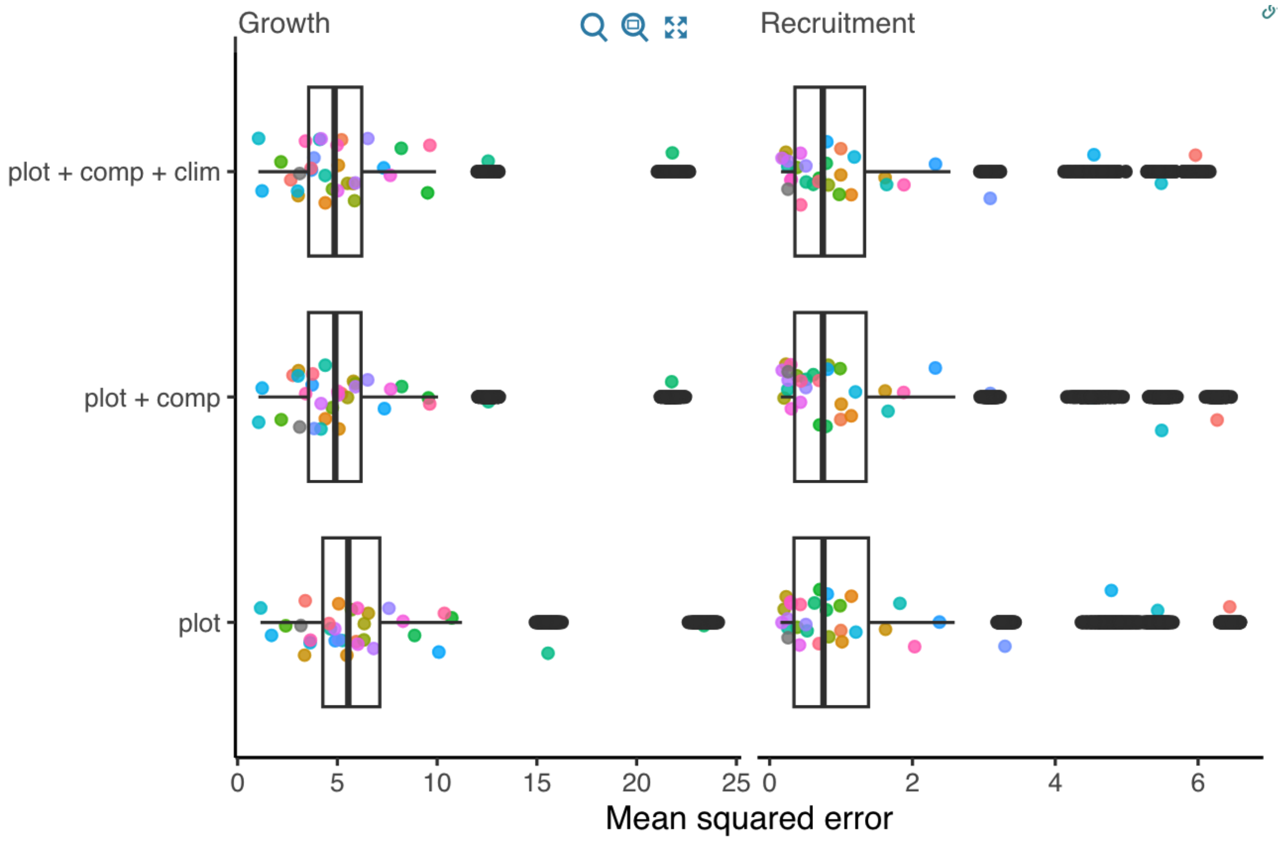

Figure 3: Posterior distribution of Mean Squared Error (MSE) across the 31 tree species as models become more complex.

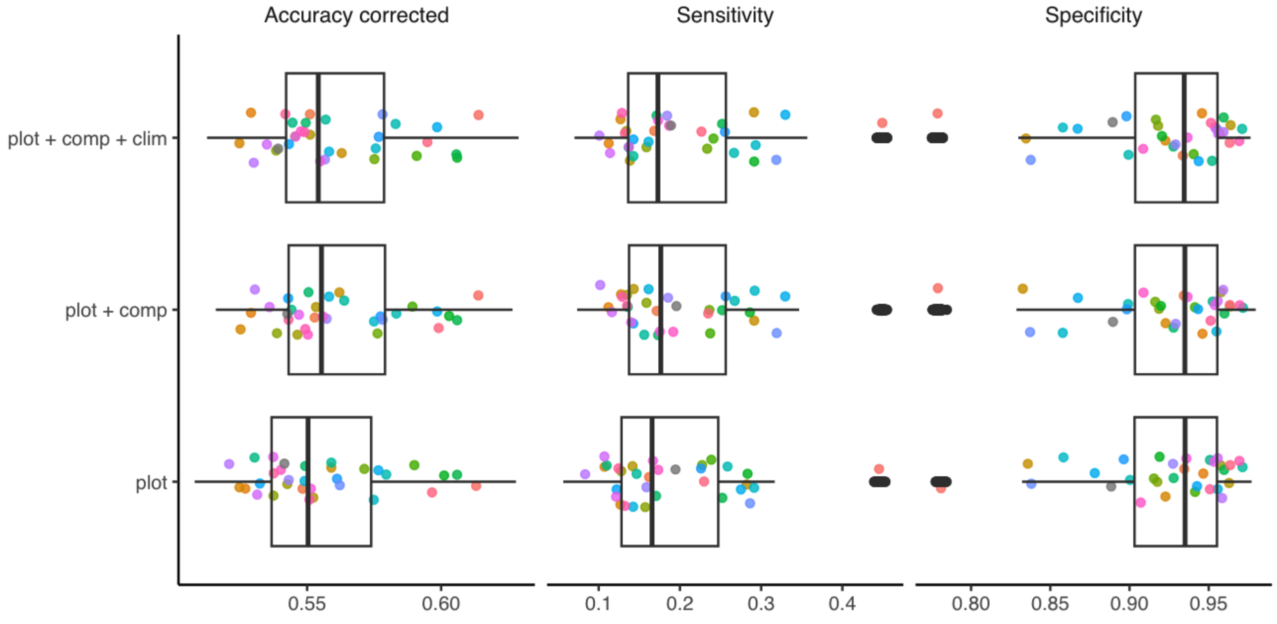

Figure 4: Comparing the posterior distribution of sensitivity, specificity, and accuracy across the 31 tree species between the competing models. Each colored dot represents the species' average posterior distribution.

the reference model in pointwise predictive cross-validation. Given the large number of observations in the dataset, we approximated LOO-CV using PSIS-LOO and subsampling. For each species, we approximated LOO-CV by sampling one-fifth of the total number of observations.

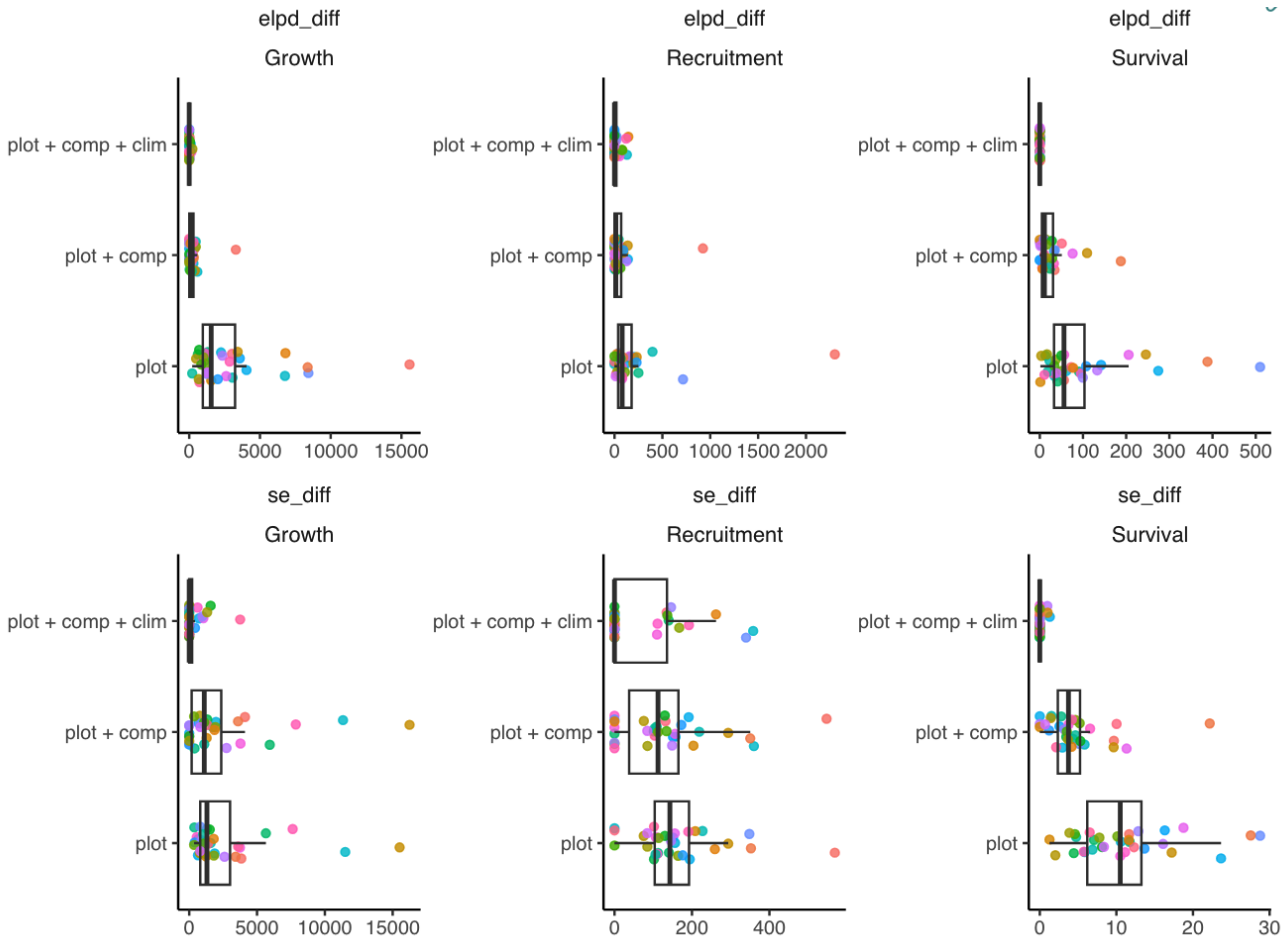

Figure 5: Boxplot shows the LOO-CV compare between the competing models based on the expected log pointwise predictive density (ELPD_diff) difference across the 31 tree species. The sd_diff is the standard error of the ELPD difference between the model and the reference model (ELPD_diff equal to zero).

#### 1.6 Size effect in survival

We initially incorporated the size effect into the survival models due to the structured-population approach. However, we observed that the effect of size on mortality probability was generally weak and variable among species, with no clear pattern of increased mortality probability with larger individual size. All models that included the size effect performed worse than the null model, which contained only plot random effects (Figure 6).

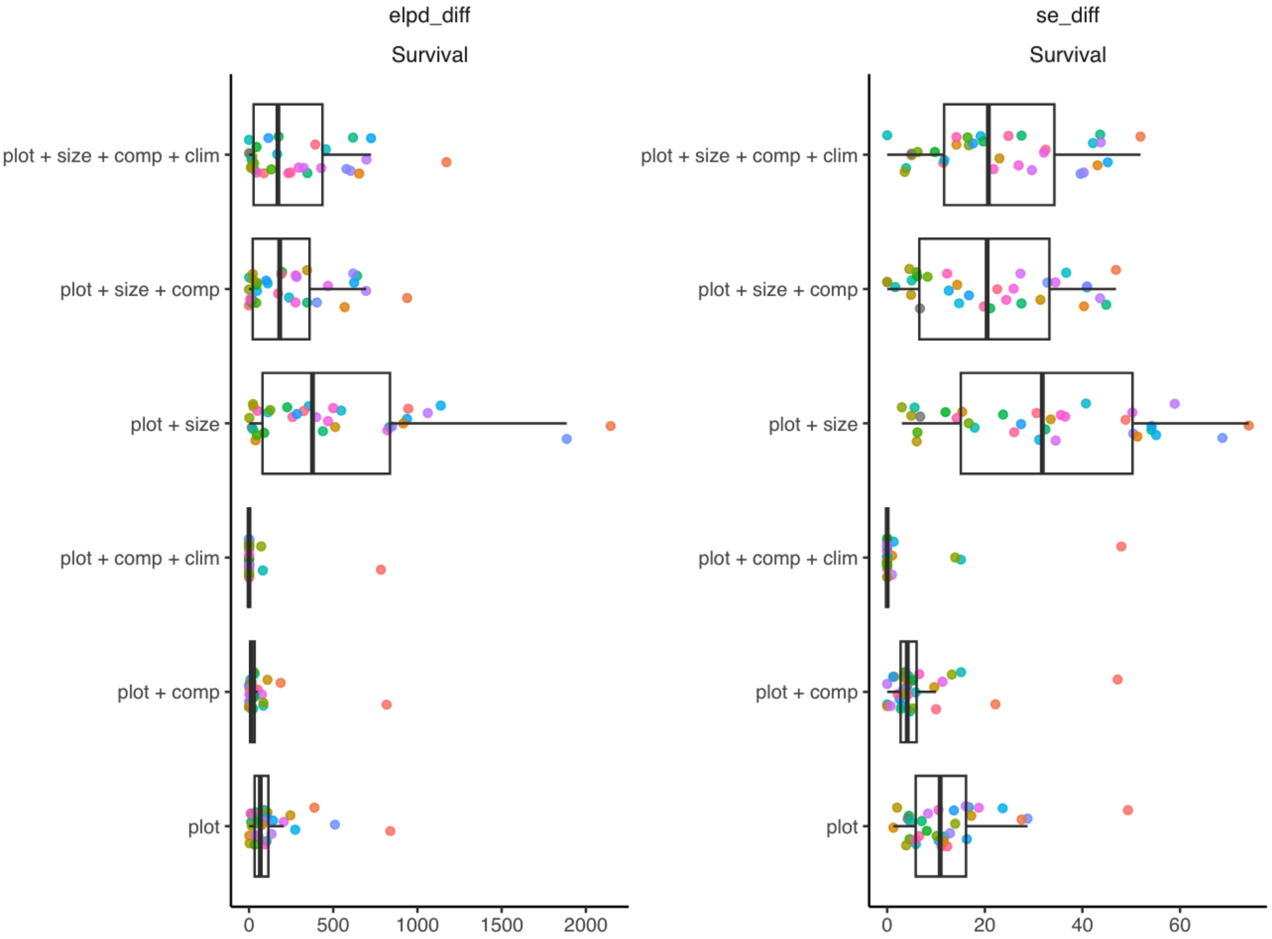

Figure 6: Boxplot shows the LOO-CV compare between the competing models based on the expected log pointwise predictive density (ELPD_diff) difference across the 31 tree species. The sd_diff is the standard error of the ELPD difference between the model and the reference model (ELPD_diff equal to zero).

#### 1.7 Conclusion

Our analysis revealed that incorporating competition into the growth, survival, and recruitment models proved more effective in gaining individual-level information than climate variables. The parameter $\sigma_{plot}$, interpreted as spatial heterogeneity, was lowest in the growth model, followed by recruitment and survival. As the models became more complex with the inclusion of covariates, recruitment exhibited the most significant reduction in spatial variance, followed by growth, with no clear pattern in the case of survival.

Regarding predictive performance, competition contributed more to the overall predictive capacity ($R^2$, MSE, and corrected accuracy) in the growth and survival models compared to climate variables. Although recruitment had the largest reduction in $\sigma_{plot}$, it had minimal impact on prediction accuracy.

Finally, the LOO-CV indicates a clear trend where the complete model featuring plot random effects, competition, and climate covariates outperformed the other competing models. Furthermore, the absolute value of the ELPD shows that the growth model gained the most information from including covariates, followed by recruitment and survival models. Consequently, we selected the complete model with plot random effects, competition, and climate covariates as the preferred model for further analysis.

### 2 Supplementary Material 2

Table 1: List of species and their frequency across the dataset.

| Species | Number of plots | Number of individual | Number of observation |
|---|---|---|---|
| *Acer rubrum* | 13149 | 96739 | 235408 |
| *Abies balsamea* | 11932 | 247737 | 521565 |
| *Betula papyrifera* | 9508 | 78049 | 203500 |
| *Picea mariana* | 7869 | 186491 | 454246 |
| *Acer saccharum* | 7403 | 71961 | 184641 |
| *Picea glauca* | 5889 | 27641 | 65626 |
| *Populus tremuloides* | 5876 | 56010 | 127115 |
| *Betula alleghaniensis* | 5624 | 28872 | 73116 |
| *Quercus rubra* | 4549 | 18272 | 46341 |
| *Quercus alba* | 4200 | 20376 | 51466 |
| *Fagus grandifolia* | 3819 | 21784 | 51764 |
| *Prunus serotina* | 3730 | 12178 | 26464 |
| *Thuja occidentalis* | 3230 | 51312 | 125811 |
| *Pinus strobus* | 3165 | 15638 | 38470 |
| *Fraxinus americana* | 2885 | 8942 | 21501 |
| *Quercus velutina* | 2722 | 10068 | 23298 |
| *Tsuga canadensis* | 2604 | 17914 | 45198 |
| *Nyssa sylvatica* | 2436 | 6275 | 15785 |
| *Quercus stellata* | 2279 | 14707 | 32102 |
| *Picea rubens* | 2190 | 16580 | 41674 |
| *Liquidambar styraciflua* | 2154 | 11655 | 29671 |
| *Fraxinus pennsylvanica* | 2149 | 9048 | 20588 |
| *Tilia americana* | 2059 | 8415 | 21412 |
| *Pinus banksiana* | 2057 | 34122 | 75372 |
| *Populus grandidentata* | 2015 | 13759 | 29358 |
| *Fraxinus nigra* | 1951 | 12633 | 31156 |

| | | | |
|---|---|---|---|
| *Liriodendron tulipifera* | 1912 | 8580 | 21071 |
| *Carya tomentosa* | 1636 | 3897 | 10590 |
| *Carya glabra* | 1622 | 4002 | 9916 |
| *Quercus prinus* | 1590 | 11000 | 27554 |
| *Juniperus virginiana* | 1571 | 9474 | 21400 |

Table 2: Prior specifications for all parameters in the species-specific demographic models. The *Symbol* column gives the notation used in the main text Methods. The *Stan name* column gives the parameter name in the publicly available Stan code. *Scale / link* indicates the transformation (if any) between the prior and the quantity referenced in the main text. Unless noted, normal priors with a positive lower bound are half-normal.

| Component | Symbol | Stan name | Scale / link | Prior |
|---|---|---|---|---|
| **Growth model** | | | | |
| Intercept | $\bar{\Gamma}$ | `r` | $\bar{\Gamma} = e^r$ (log-link) | $r \sim \mathcal{N}(-3.5, 1)$ |
| Asymptotic size | $\zeta_\infty$ | `Lmax` | cm; lower bound = 1.2 × max(DBH) | $\zeta_\infty \sim \mathcal{N}(1000, 80)$ |
| Observation s.d. | $\sigma$ | `sigma_obs` | half-normal | $\sigma \sim \mathcal{N}^+(0, 1.5)$ |
| Plot effect | $\alpha_j$ | `rPlot_log` | on log-$\Gamma$ scale | $\alpha_j \sim \mathcal{N}(0, \sigma_\alpha)$ |
| Plot s.d. | $\sigma_\alpha$ | `sigma_plot` | half-normal | $\sigma_\alpha \sim \text{Exp}(2)$ |
| Competition | $\beta$ | `Beta` | on log-$\Gamma$ scale | $\beta \sim \mathcal{N}(-1, 1)$ |
| Comp. partition | $\theta$ | `theta` | lower bound = 0 | $\theta \sim \text{LogNormal}(1, 3)$ |
| Optimal MAT | $\xi_{\text{MAT}}$ | `optimal_temp` | scaled to $[0, 1]$ within species range | $\xi_{\text{MAT}} \sim \text{Beta}(2, 2)$ |
| MAT breadth | $\sigma_{\text{MAT}}$ | `tau_temp` | $\tau_{\text{MAT}} = 1/\sigma^2_{\text{MAT}}$; half-normal | $\tau_{\text{MAT}} \sim \mathcal{N}^+(0, 1)$ |
| Optimal MAP | $\xi_{\text{MAP}}$ | `optimal_prec` | scaled to $[0, 1]$ within species range | $\xi_{\text{MAP}} \sim \text{Beta}(2, 2)$ |

*(continued on next page)*

(continued from previous page)

| Component | Symbol | Stan name | Scale / link | Prior |
|---|---|---|---|---|
| MAP breadth | $\sigma_{\text{MAP}}$ | `tau_prec` | $\tau_{\text{MAP}} = 1/\sigma^2_{\text{MAP}}$; half-normal | $\tau_{\text{MAP}} \sim \mathcal{N}^+(0,\,1)$ |
| **Survival model** | | | | |
| Intercept | $\bar{\psi}$ | `psi` | $\bar{\psi} = \text{logit}^{-1}(\texttt{psi})$; bounds $[-2, 10]$ | $\texttt{psi} \sim \mathcal{N}(5,\,1)$ |
| Plot effect | $\alpha_j$ | `psiPlot` | on logit-$\psi$ scale | $\alpha_j \sim \mathcal{N}(0,\,\sigma_\alpha)$ |
| Plot s.d. | $\sigma_\alpha$ | `sigma_plot` | lognormal | $\sigma_\alpha \sim$ LogNormal(2, 1) |
| Competition | $\beta$ | `Beta` | on logit-$\psi$ scale | $\beta \sim \mathcal{N}(0,\,0.1)$ |
| Comp. partition | $\theta$ | `theta` | bounds $[0, 2]$ | $\theta \sim$ Exp(2.5) |
| Optimal MAT | $\xi_{\text{MAT}}$ | `optimal_temp` | scaled to $[0, 1]$ within species range | $\xi_{\text{MAT}} \sim$ Beta(2, 2) |
| MAT breadth | $\sigma_{\text{MAT}}$ | `tau_temp` | $\tau_{\text{MAT}} = 1/\sigma^2_{\text{MAT}}$; half-normal | $\tau_{\text{MAT}} \sim \mathcal{N}^+(0,\,2)$ |
| Optimal MAP | $\xi_{\text{MAP}}$ | `optimal_prec` | scaled to $[0, 1]$ within species range | $\xi_{\text{MAP}} \sim$ Beta(2, 2) |
| MAP breadth | $\sigma_{\text{MAP}}$ | `tau_prec` | $\tau_{\text{MAP}} = 1/\sigma^2_{\text{MAP}}$; half-normal | $\tau_{\text{MAP}} \sim \mathcal{N}^+(0,\,2)$ |
| **Recruitment model** | | | | |
| Ingrowth rate | $\bar{\phi}$ | `mPop_log` | $\bar{\phi} = e^{\texttt{mPop\_log}}$ (log-link) | $\texttt{mPop\_log} \sim \mathcal{N}(-5,\,1.5)$ |
| Plot effect | $\alpha_j$ | `mPlot_log` | on log-$\phi$ scale | $\alpha_j \sim \mathcal{N}(0,\,\sigma_\alpha)$ |
| Plot s.d. | $\sigma_\alpha$ | `sigma_plot` | half-normal | $\sigma_\alpha \sim$ Exp(6) |
| Optimal density | $\delta$ | `optimal_BA` | $\text{m}^2\,\text{ha}^{-1}$; bounds $[5, 150]$ | $\delta \sim \mathcal{N}(20,\,10)$ |

(continued on next page)

*(continued from previous page)*

| Component | Symbol | Stan name | Scale / link | Prior |
|---|---|---|---|---|
| Density breadth | $\sigma$ | `sigma_BA` | half-normal | $\sigma \sim \mathcal{N}^+(15,\, 10)$ |
| Recruit survival | $\bar{\rho}$ | `p_log` | $\bar{\rho} = \exp(-e^{\texttt{p\_log}})$ | $\texttt{p\_log} \sim \mathcal{N}(-3,\, 1.5)$ |
| Comp. on $\rho$ | $\beta_p$ | `beta_p` | half-normal; $\theta \equiv 1$ for $\rho$ | $\beta_p \sim \mathcal{N}^+(0,\, 0.6)$ |
| Optimal MAT | $\xi_{\mathrm{MAT}}$ | `optimal_temp` | scaled to $[0, 1]$ within species range | $\xi_{\mathrm{MAT}} \sim \mathrm{Beta}(2,\, 2)$ |
| MAT breadth | $\sigma_{\mathrm{MAT}}$ | `tau_temp` | $\tau_{\mathrm{MAT}} = 1/\sigma^2_{\mathrm{MAT}}$; half-normal | $\tau_{\mathrm{MAT}} \sim \mathcal{N}^+(0,\, 2)$ |
| Optimal MAP | $\xi_{\mathrm{MAP}}$ | `optimal_prec` | scaled to $[0, 1]$ within species range | $\xi_{\mathrm{MAP}} \sim \mathrm{Beta}(2,\, 2)$ |
| MAP breadth | $\sigma_{\mathrm{MAP}}$ | `tau_prec` | $\tau_{\mathrm{MAP}} = 1/\sigma^2_{\mathrm{MAP}}$; half-normal | $\tau_{\mathrm{MAP}} \sim \mathcal{N}^+(0,\, 2)$ |
| **Recruit size model** | | | | |
| Size intercept | $\Omega$ | `size_int` | cm; half-normal, $L_{\mathrm{rec}} = 12.7$ | $\Omega \sim \mathcal{N}^+(L_{\mathrm{rec}},\, 50)$ |
| Time slope | $\gamma_{\mathrm{rec}}$ | `phi_time` | $\mathrm{cm\,yr}^{-1}$; half-normal | $\gamma_{\mathrm{rec}} \sim \mathcal{N}^+(0,\, 1)$ |
| Size s.d. | $\sigma$ | `sigma_size` | half-normal | $\sigma \sim \mathcal{N}^+(20,\, 5)$ |

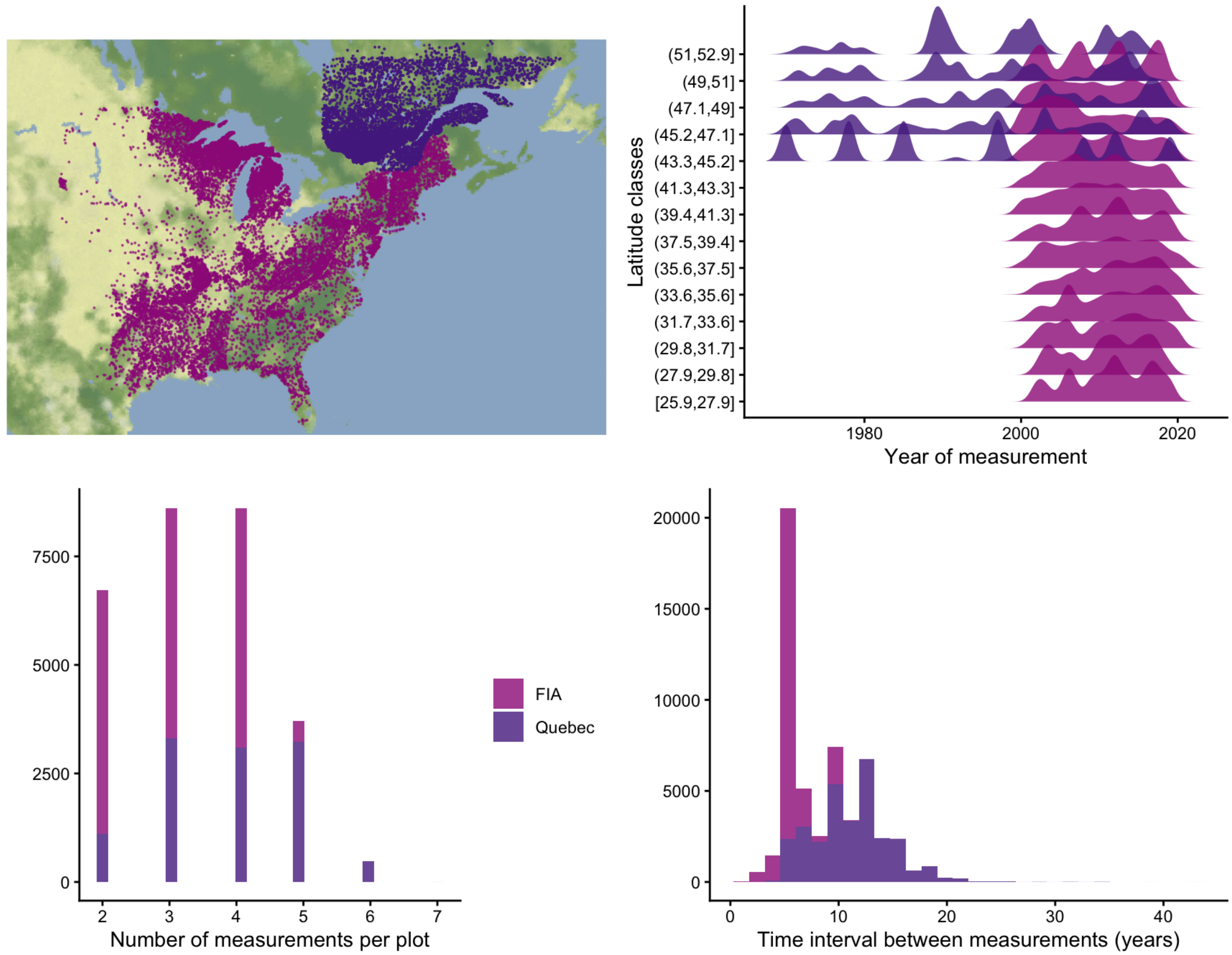

Figure 7: Spatial (top left) and temporal (top right) coverage of the dataset incorporating data from the USA and Quebec. The top right panel shows the distribution of observations per class of latitude for the 31 species used in this study.

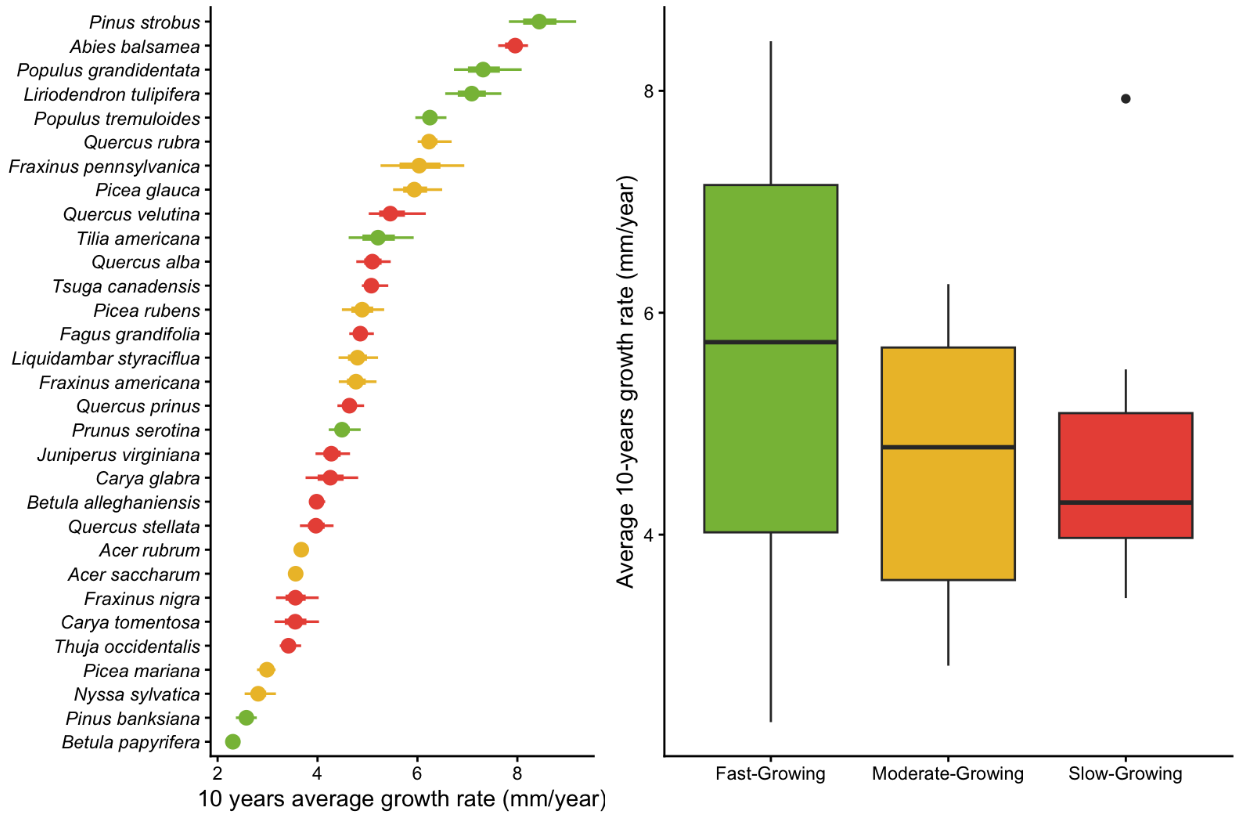

Figure 8: Posterior distribution for the intercept of the growth model using the 10-year average growth rate. Species are classified by their general growth trait following Burns et al. (1990).

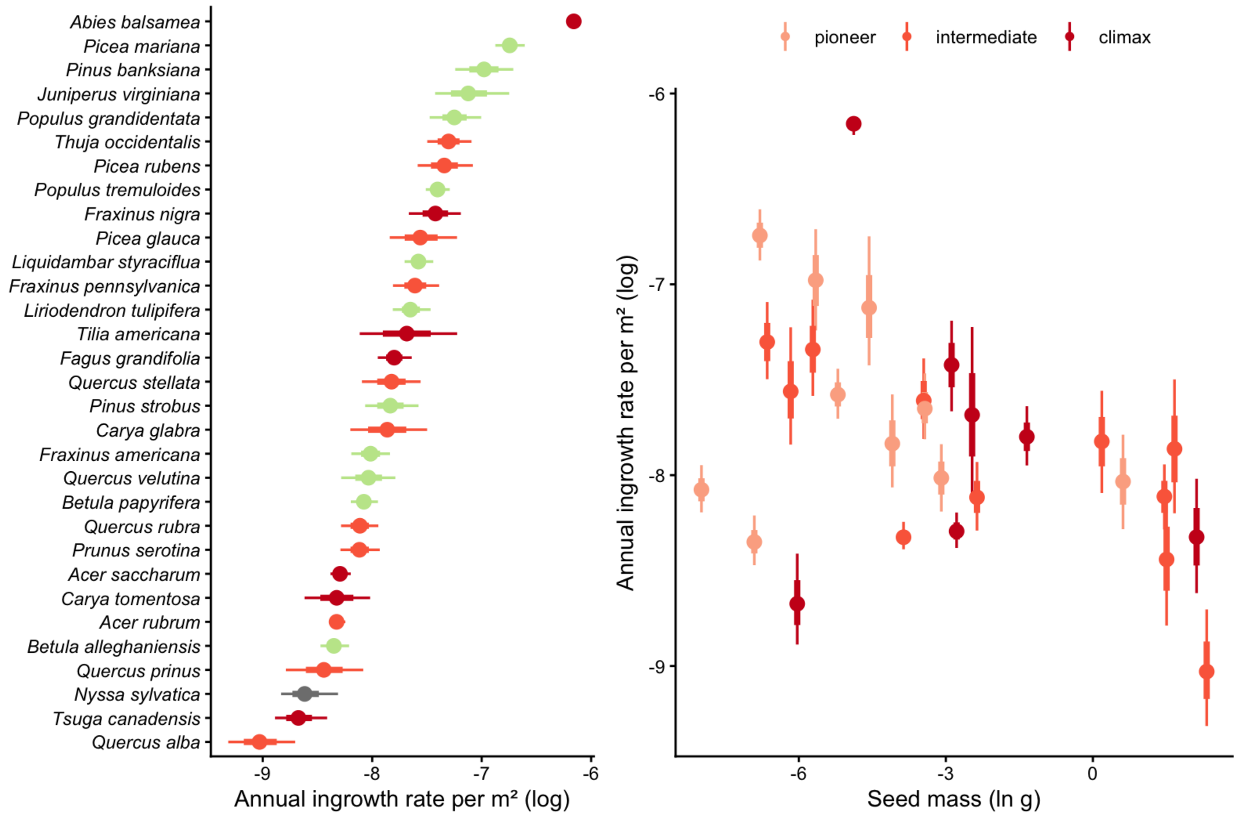

Figure 9: Posterior distribution for the intercept of the ingrowth model for the number of individuals that ingress the population per year per $m^2$ in function seed mass (Díaz et al. 2022). Species are classified by their successional status following Burns et al. (1990).

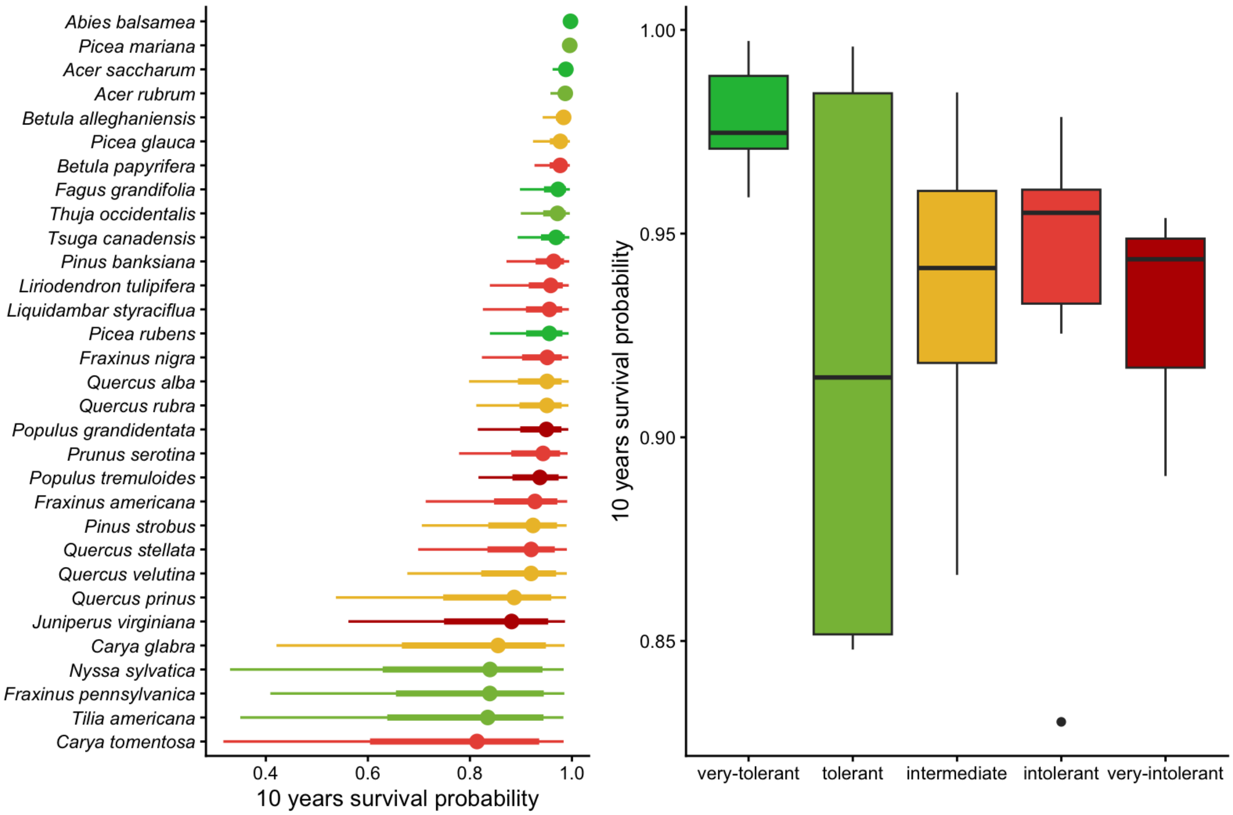

Figure 10: Posterior distribution for the intercept of the annual survival probability for the ingrowth model. Species are classified by their shade tolerance trait following Burns et al. (1990).

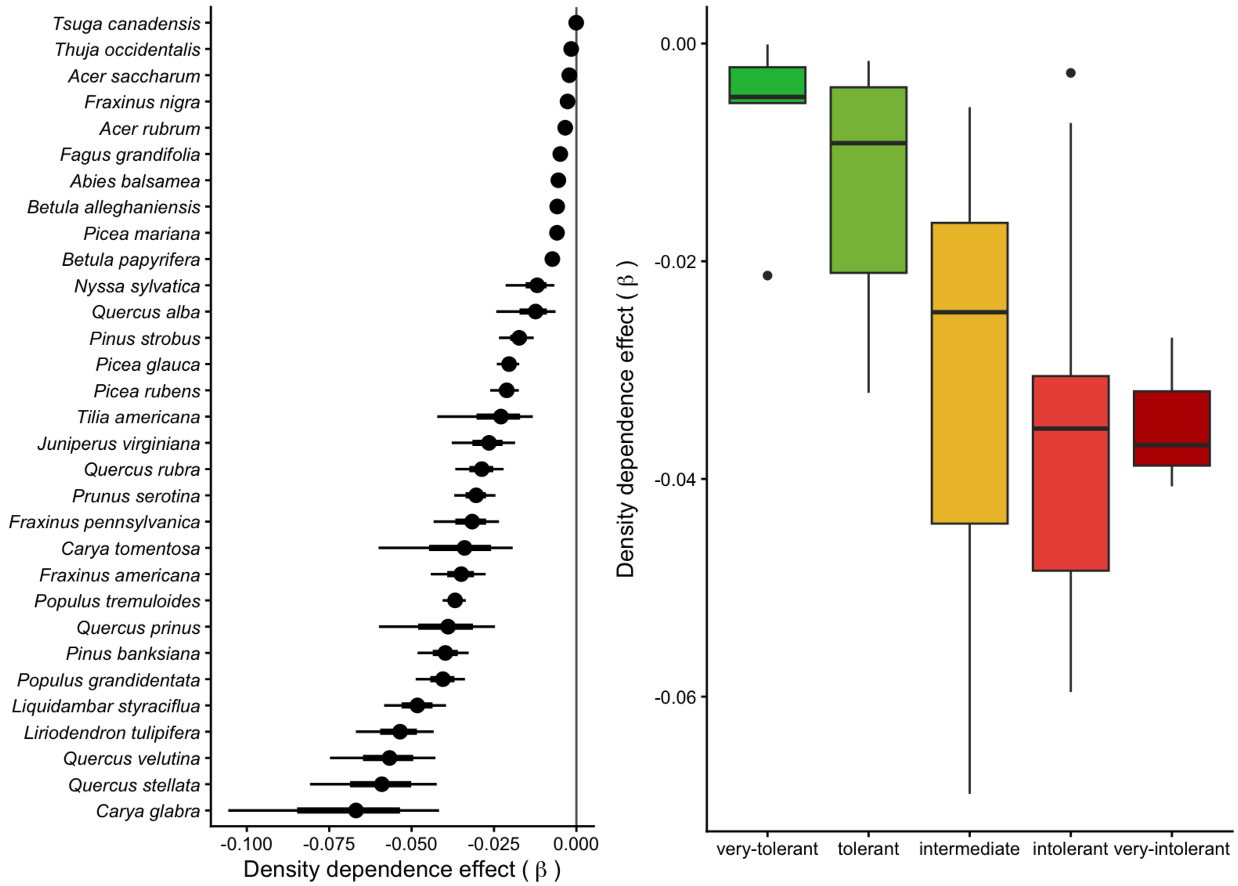

Figure 11: Posterior distribution of the density dependence parameter affecting the annual survival rate of recruitment individuals. Species are classified by their shade tolerance trait following Burns et al. (1990).

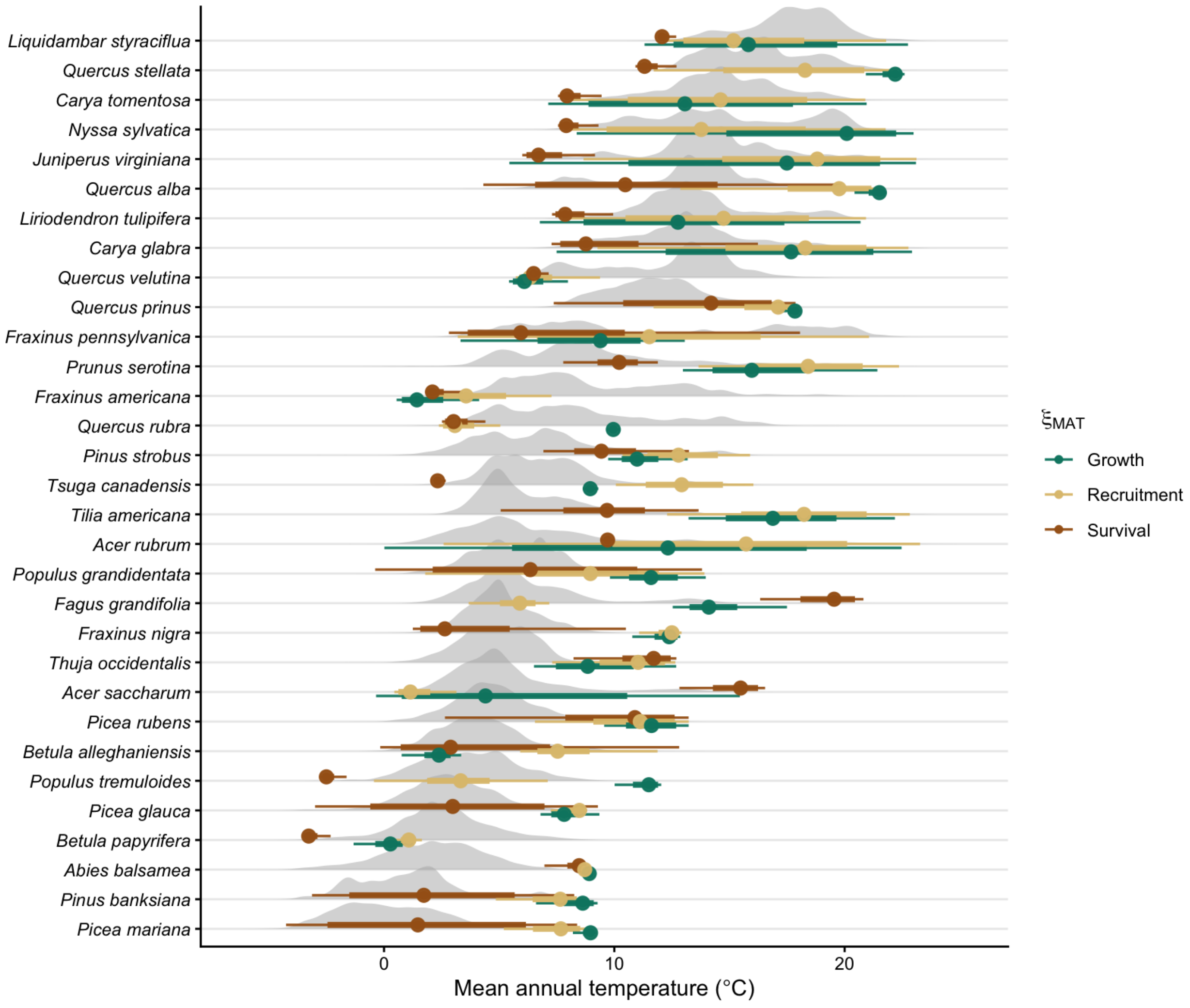

Figure 12: Distribution for the optimal annual mean temperature ($\xi_{MAT}$) for growth (green), recruitment (yellow), and survival (brown). The gray density plot is the annual mean temperature distribution among all observed trees across space and time.

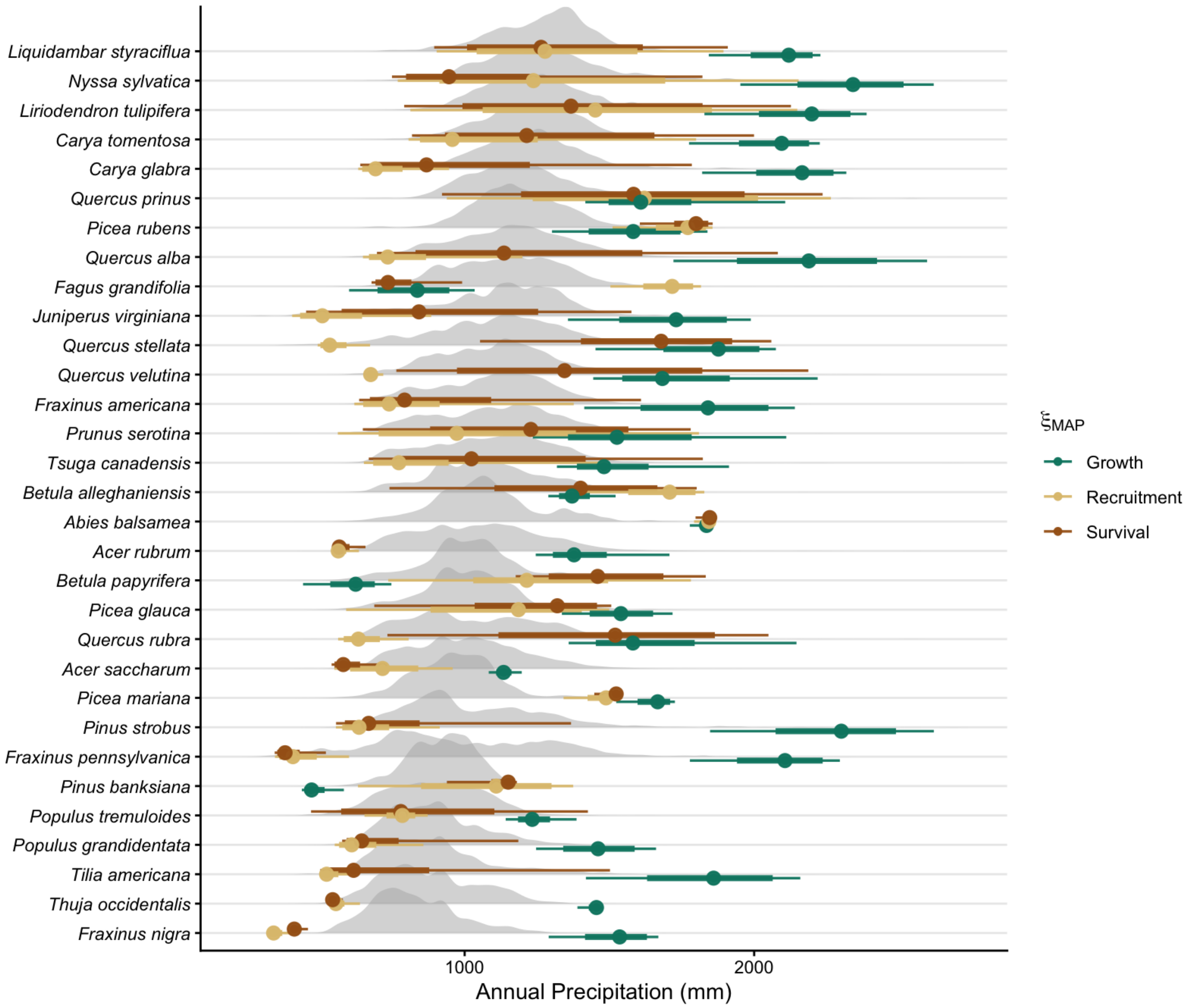

Figure 13: Distribution for the optimal mean annual precipitation ($\xi_{MAP}$) for growth (green), recruitment (yellow), and survival (brown). The density plot in gray is the distribution of annual precipitation variable among all observed trees across space and time.

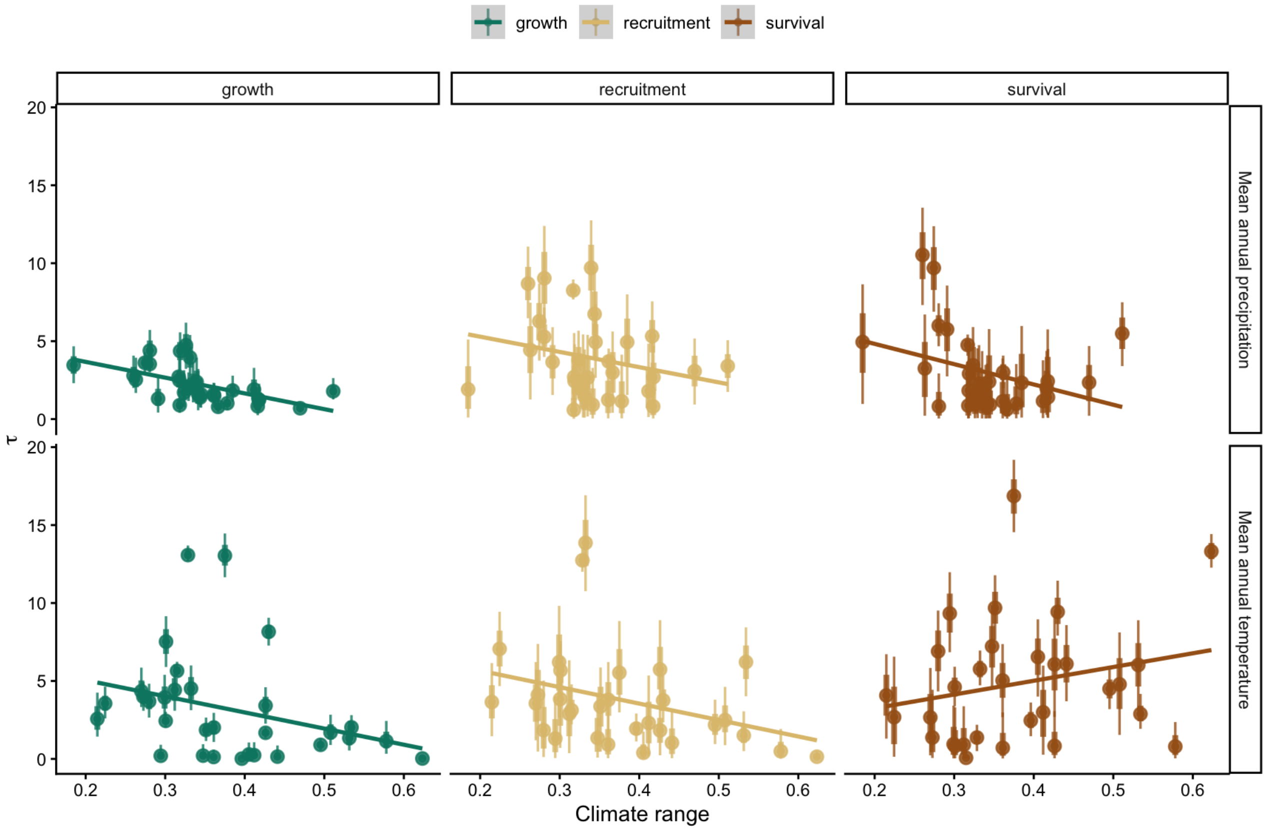

Figure 14: Climate breadth in function of climate range size. The higher the climate range size, the more climate conditions the species experienced.

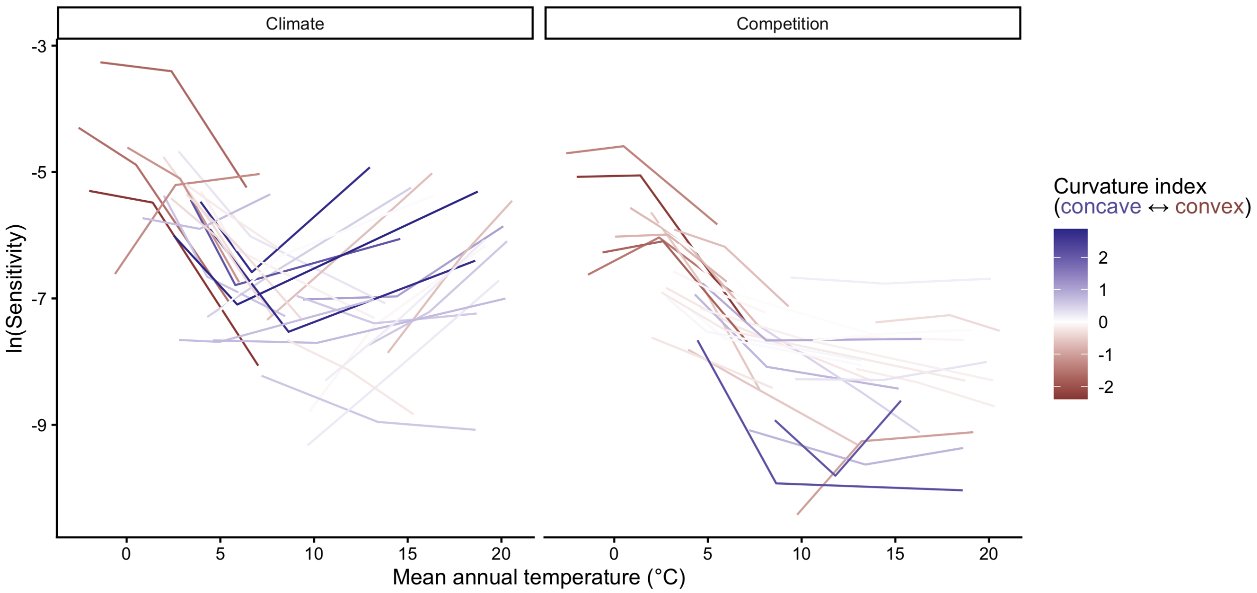

Figure 15: Sensitivity of species population growth rate to climate (left) and competition (right). Each species is represented by a connected line linking their cold, center, and hot range position points. Each of these 3 values represents the average sensitivity across all plots classified as one these three range position groups. Each line was colored according to a curvature index, ranging from convace (blue), linear (transparent), to convex (red). Range positions were defined using the median mean annual temperature (MAT) across all plots belonging to each thermal class. Note that uncertainty in each sensitivity point estimation has been omitted for clarity.

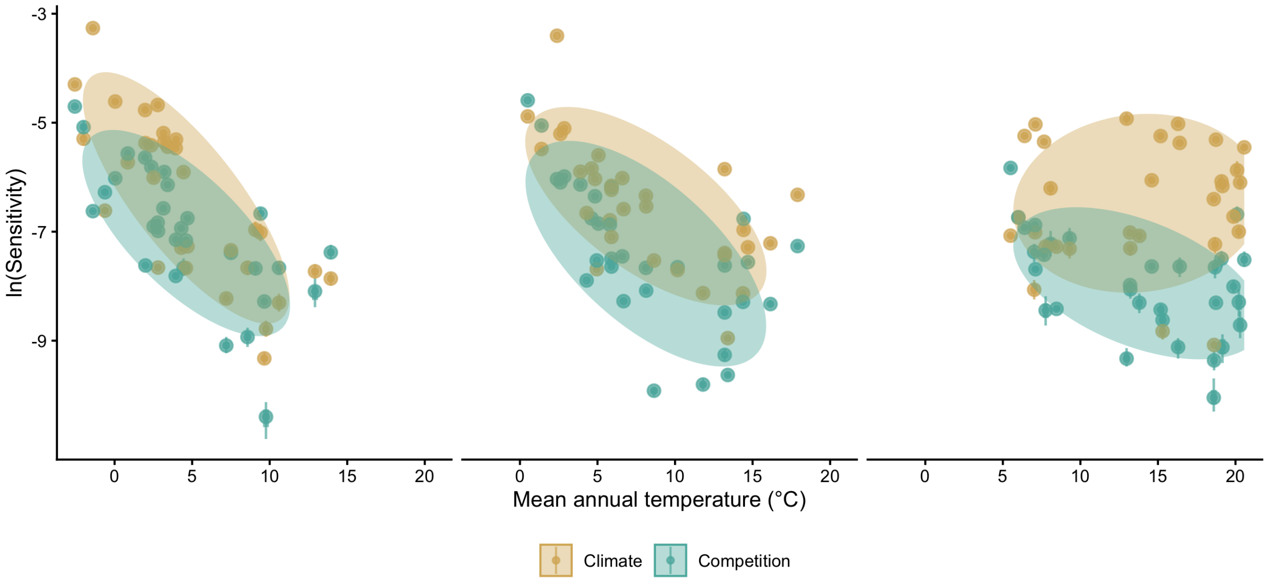

Figure 16: Sensitivity of species population growth rate to competition (green) and climate (yellow) across the cold, center, and hot temperature ranges. Range positions were defined using the median mean annual temperature (MAT) across all plots belonging to each thermal class. In the bottom panel, species points are grouped by a Multivariate Normal Density function with 75% probability, while in the top panel, the lines represent the 25, 50, and 75% quantile probabilities.

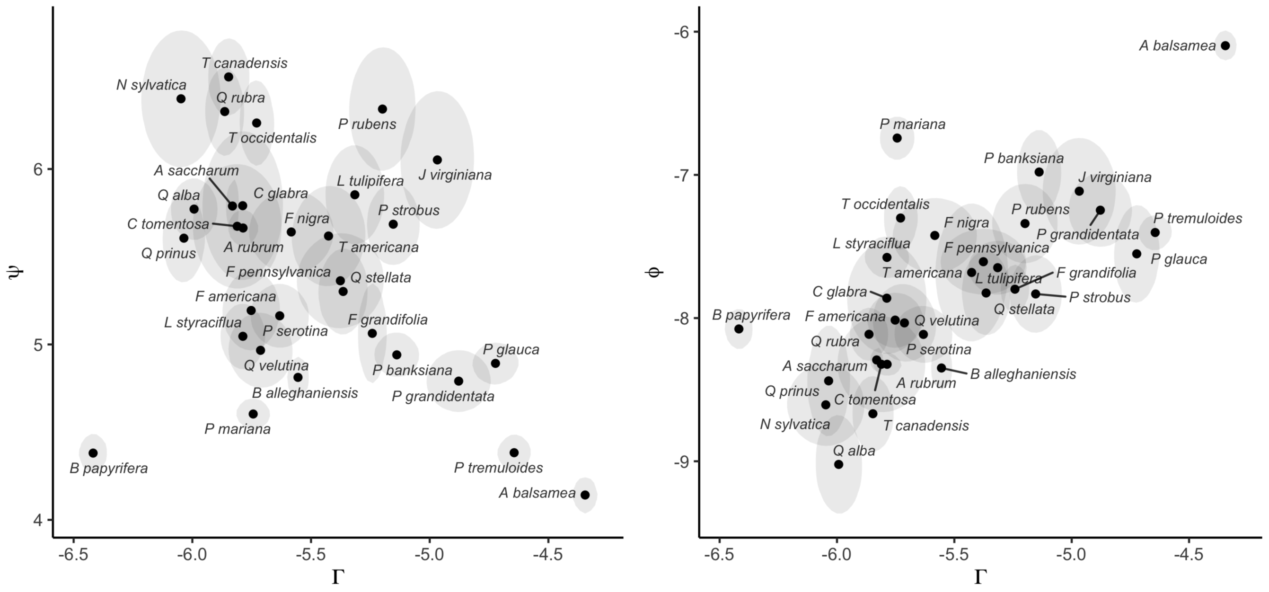

Figure 17: Correlation between (left panel) growth rate ($\Gamma$) and annual survival rate ($\psi$) and (right panel) growth rate ($\Gamma$) and annual recruitment rate ($\phi$). The uncertainty of the parameters is summarised by a Multivariate Normal Density function with 90% probability.

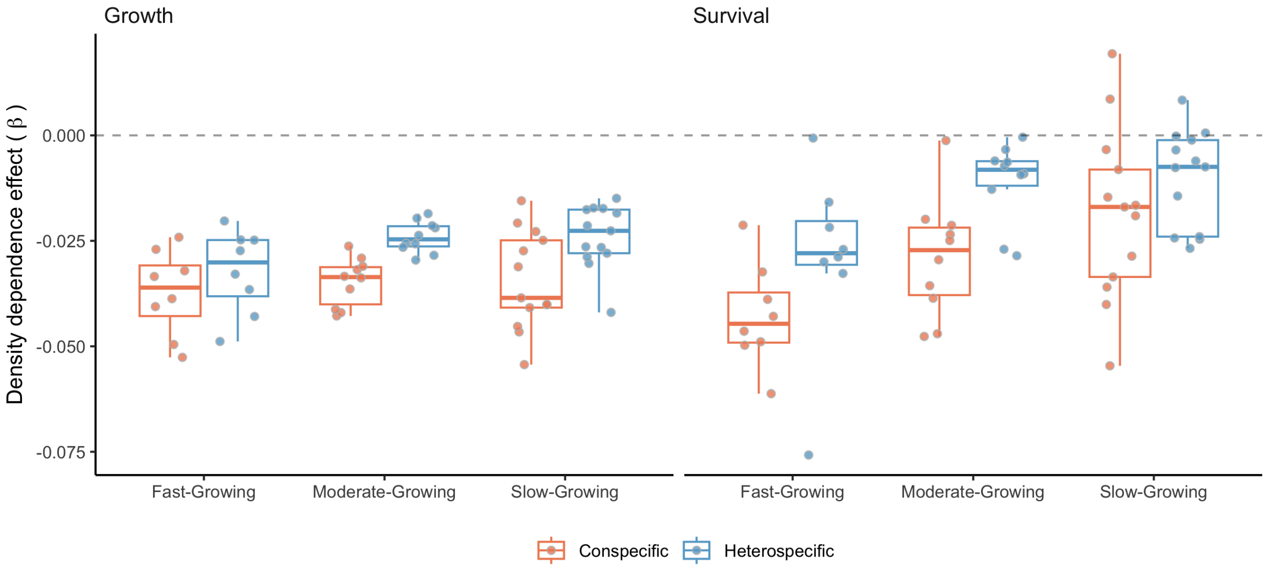

Figure 18: Posterior distribution for the conspecific (red) and heterospecific (blue) density dependence for each class of growth rate (Burns et al. 1990). The more negative the $\beta$, the stronger the competition effect.

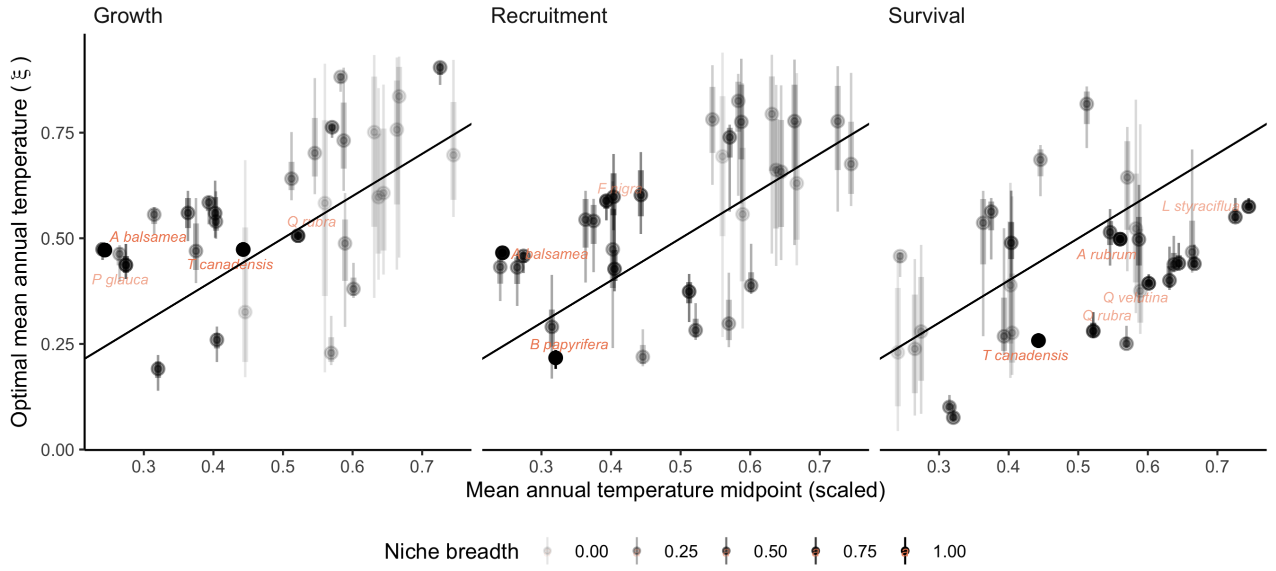

Figure 19: Correlation between posterior distribution of optimal temperature ($\xi_{MAT}$) and the species' midpoint location across the mean annual temperature range. The transparency of each species point is scaled to be a function of niche breadth. The closer this value is to zero, the higher the breadth around the mean. In other words, when climate breadth is zero, the bell-shaped unimodal function becomes an almost flat line. Colored species names are those with niche breadth higher than 0.5.

### 3 Supplementary Material 3

#### 3.1 Sensitivity analysis

Here, we conducted a global sensitivity analysis (GSA) of the population growth rate ($\lambda$) with respect to demographic models. Sensitivity analysis uses various methods to decompose the total variance of an outcome into contributions from parameters or input variables. In structured population models, sensitivity analyses involve computing partial derivatives of $\lambda$ to individual parameters, following Caswell (1978) as:

$$\frac{\partial \lambda}{\partial \theta_i} \tag{1}$$

where theta represents a vector of $i$ parameters. However, most methods quantify the local sensitivity of each parameter separately while holding all others constant (Saltelli et al. 2019). This approach can overlook the obscure parameter interactions often common in complex models. Furthermore, because of the high dimensionality of IPM due to the large number of parameters, these methods can quickly become computationally expensive.

To address this, we leveraged the efficiency of non-parametric models, such as random forests, for variable importance classification (Antoniadis et al. 2021). This approach offers speed and suits our study as it allows us to quantify both sources of variability in $\lambda$. It accounts for the sensitivity of $\lambda$ to each parameter and considers the uncertainty associated with the parameters. Therefore, a specific parameter may have higher importance because either $\lambda$ is more sensitive to it or because the parameter is more uncertain.

We quantified the variability in population growth rate in function of the parameters using an *insileco* experimental approach. Specifically, we quantified the variability $\lambda$ for different climate conditions, ranging from cold to the center and up to the hot mean annual temperatures experienced by each species. Furthermore, we combined the climate conditions with a low and high competition intensity. We defined the temperature ranges for each species using the 1st, 50th, and 99th percentiles. The low competition was defined as a population size of $N = 0.1$, while high competition was set at the 99th percentile of the plot basal area. Precipitation was kept at optimal conditions computed based on the average optimal precipitation parameters among growth, survival, and recruitment models.

For each species, climate, and competition conditions, we computed $\lambda$ 500 times using different draws from the posterior distribution, setting the plot random effects to zero. The code used for this analysis can be found in the `forest-IPM` GitHub repository.

##### 3.2 Simulation Summary

The final simulation involved a total of 500 draws across species and different conditions. The Figure 20 illustrates the distribution of $\lambda$ computed using 500 random draws from the posterior distribution of parameters across different climate and competition conditions.

##### 3.3 Importance of demographic models

Random forest is a non-parametric classification or regression model that ranks each input variable's importance in explaining the variance of the response variable. We used the permutation method for ranking variable importance (Breiman 2001). This method measures the change in model performance by individually shuffling (permuting) the values of each input variable. The greater the change in predictive accuracy with shuffling input values, the more important the specific variable will become. This is computed individually for each tree and then averaged across all $n$ random trees. Finally, we normalized the importance output of each regression model so that they sum to 1. We used the R package `ranger` with default hyperparameters for fitting the random forest models (Wright and Ziegler 2017).

Figure 21 shows the distribution of $R^2$ from 20 random forest replications across different climate and competition conditions. These values range from 0.2 to 0.9, with an average value of 0.63 across species and conditions. This variation possibly reflects the uncertainty in the parameters across species.

As our primary interest lies in demographic levels rather than parameter levels, we focus on the combined importance of all parameters for each demographic model. This splits the total importance among the four demographic functions of the IPM: growth, survival, recruitment, and recruited size models. The recruited size model had an insignificant contribution to $\lambda$, with nearly all random forest models showing a contribution below 1%. Thus, we omitted this model and concentrated on the growth, survival, and recruitment models, which collectively explain over 99% of the variation in $\lambda$

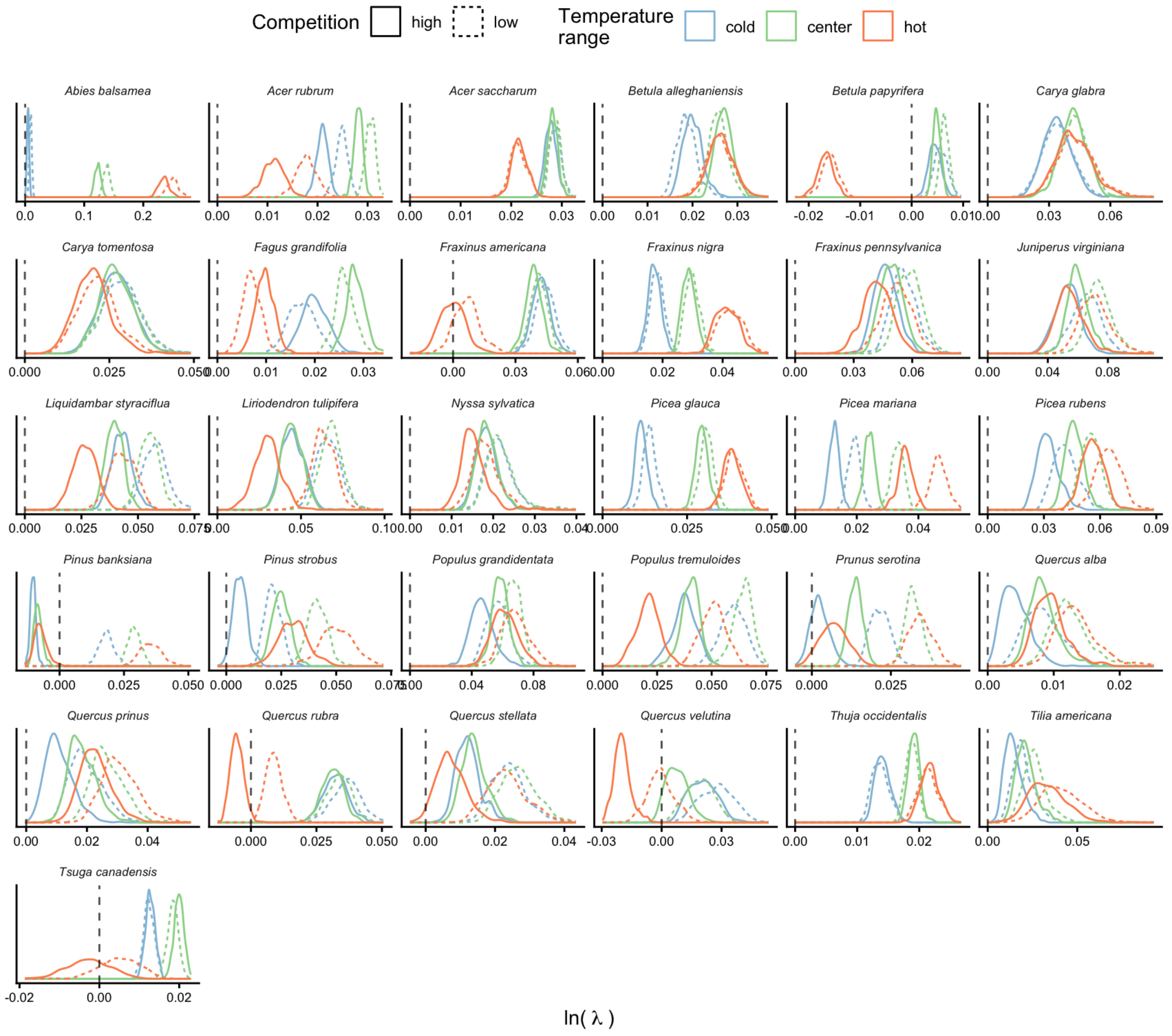

Figure 20: Distribution of 500 draws of population growth rate ($\lambda$) for different climate and competition conditions

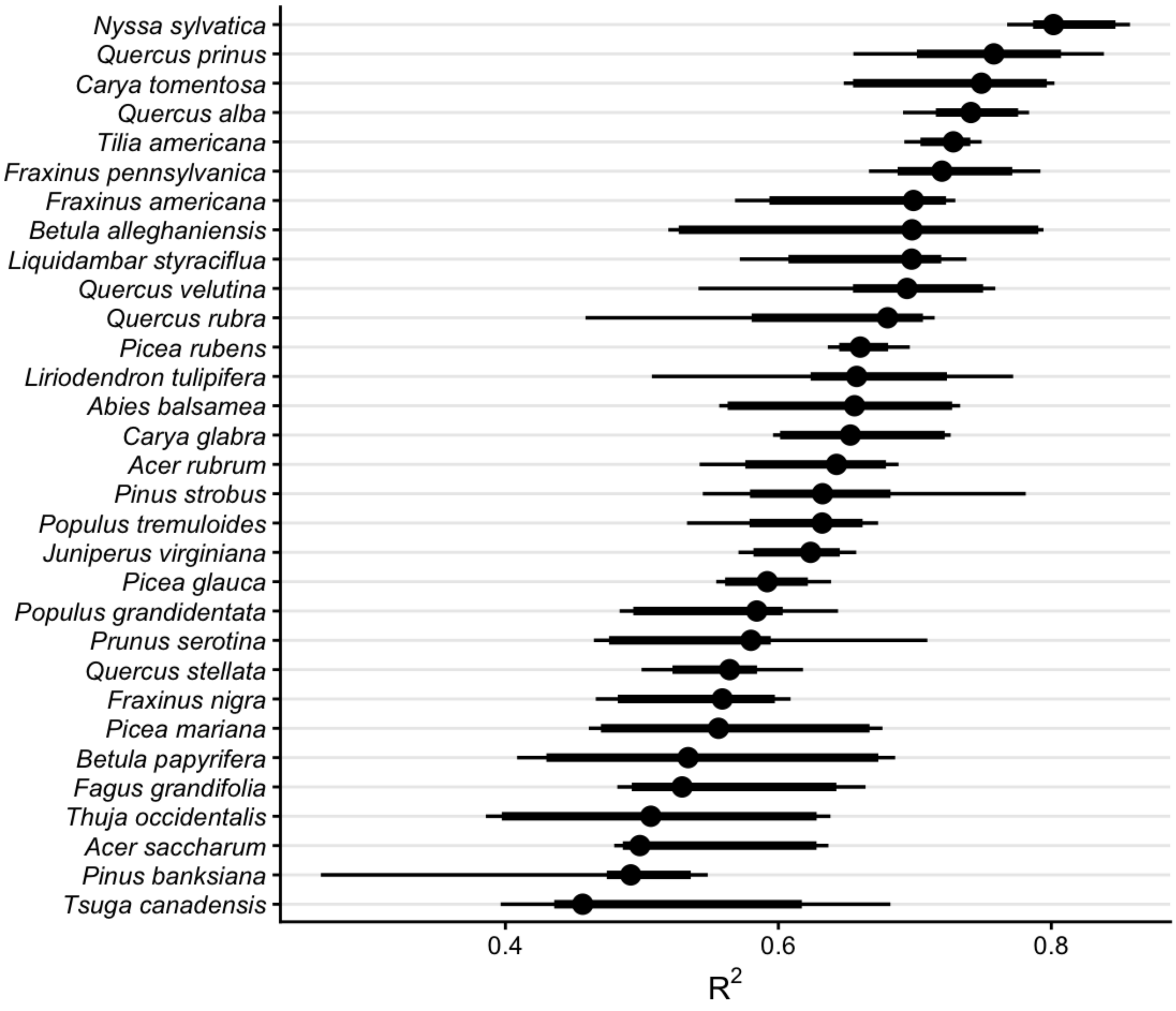

Figure 21: Distribution of $R^2$ from 20 random forest replications across different climate and competition conditions.

(Figure 22).

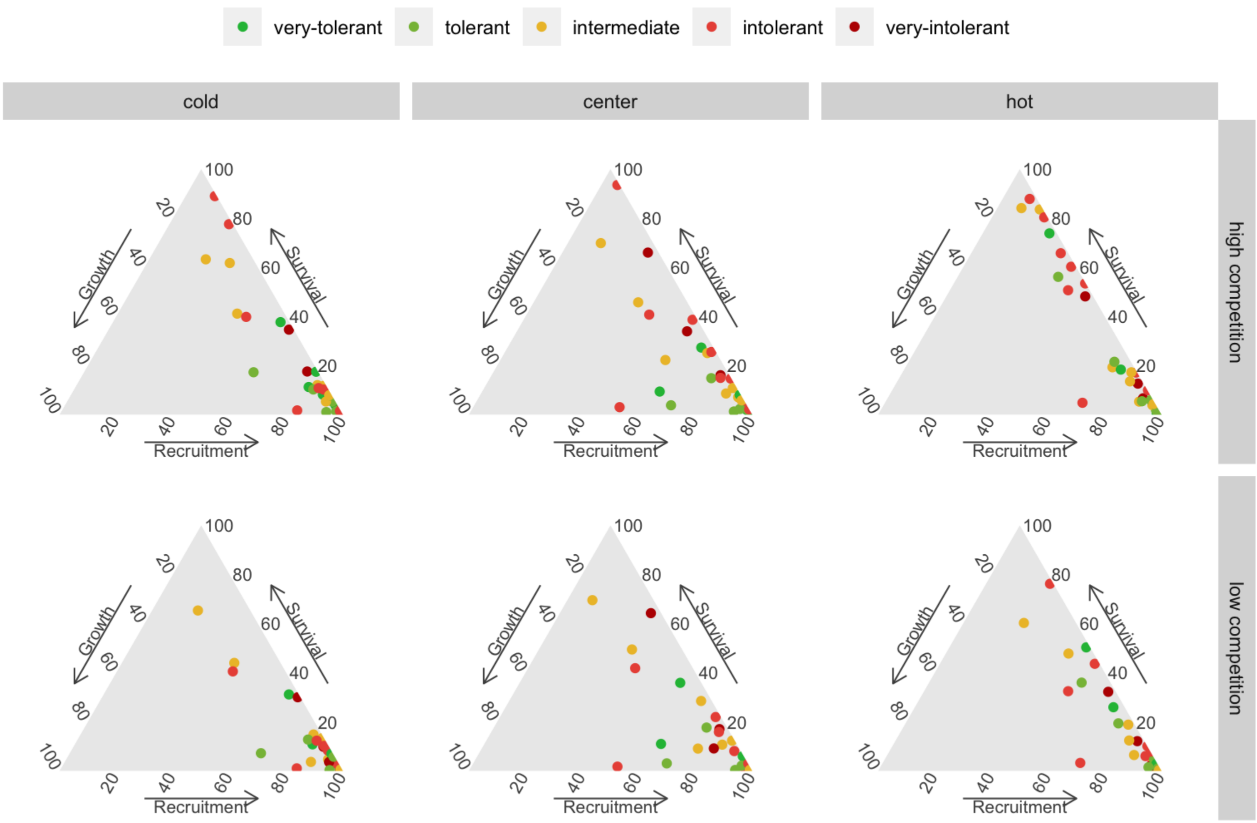

Figure 22: Ternary plot describing the importance distribution among the growth, survival, and recruitment models. Color represents the level of shade tolerance (Burns et al. 1990).

The ternary plots above show the raw importance data from the random forest, which can be challenging to interpret. The key message is that variance in $\lambda$ is primarily explained by the recruitment and survival demographic models. Furthermore, certain conditions appear to shift the importance from recruitment to the survival model. In Figure 23, we explore the correlation between the importance of recruitment and survival under different covariate conditions.

We observe that at low competition, for most species, variations in $\lambda$ are primarily explained by recruitment. This pattern slightly diminishes as we move from the cold range to the center and up to the hot temperature range. We can observe an overall shift toward the survival model at high competition intensity, especially in the hot temperature range.

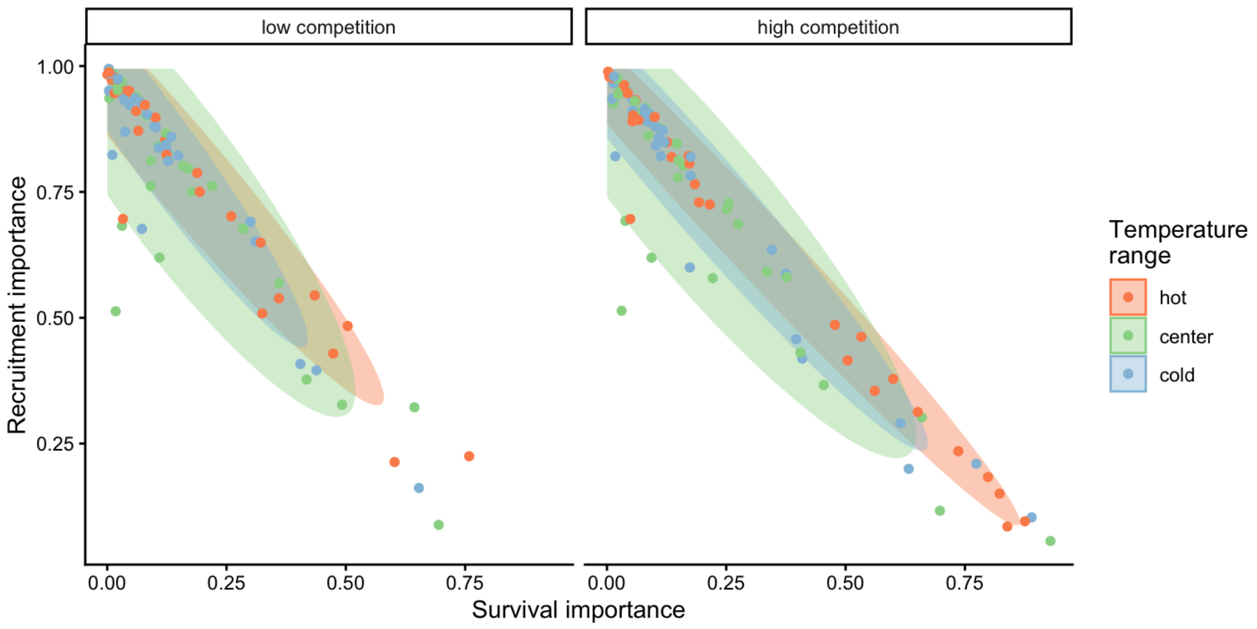

Figure 23: Correlation between the survival and recruitment relative importance across the 31 species, climate and competition conditions. Species points are grouped by a Multivariate Normal Density function with a probability of 90%.

##### 3.4 Importance of covariates

Similar to assessing parameter importance, we also used the random forest approach to evaluate the importance of covariates. For simplicity, we used the same output of the simulations as previously explained, shifting the explanatory variables from parameters to covariates.[1] The Figure 24 shows the distribution of relative importance between climate and competition covariates for each species.

##### 3.5 Notes on Conspecific and Heterospecific Competition Effects

In the preceding discussion, we did not specify whether we were considering conspecific or heterospecific competition. For all the results presented in this chapter, the *high competition* condition was applied at the heterospecific level, while conspecific competition was set to a very low proportion. This choice is based on the standard invasion growth rate metric, or the population growth rate when rare, an important measure for quantifying population persistence (Lewontin and Cohen 1969).

Additionally, we performed the sensitivity analysis with the same conditions, except for changing the high competition from heterospecific to conspecific individuals. We observed that nearly all the

[1]This analysis could be expanded to include more marginal conditions beyond just cold, center, and hot temperatures and low and high competition. However, this would exponentially increase the number of simulations.

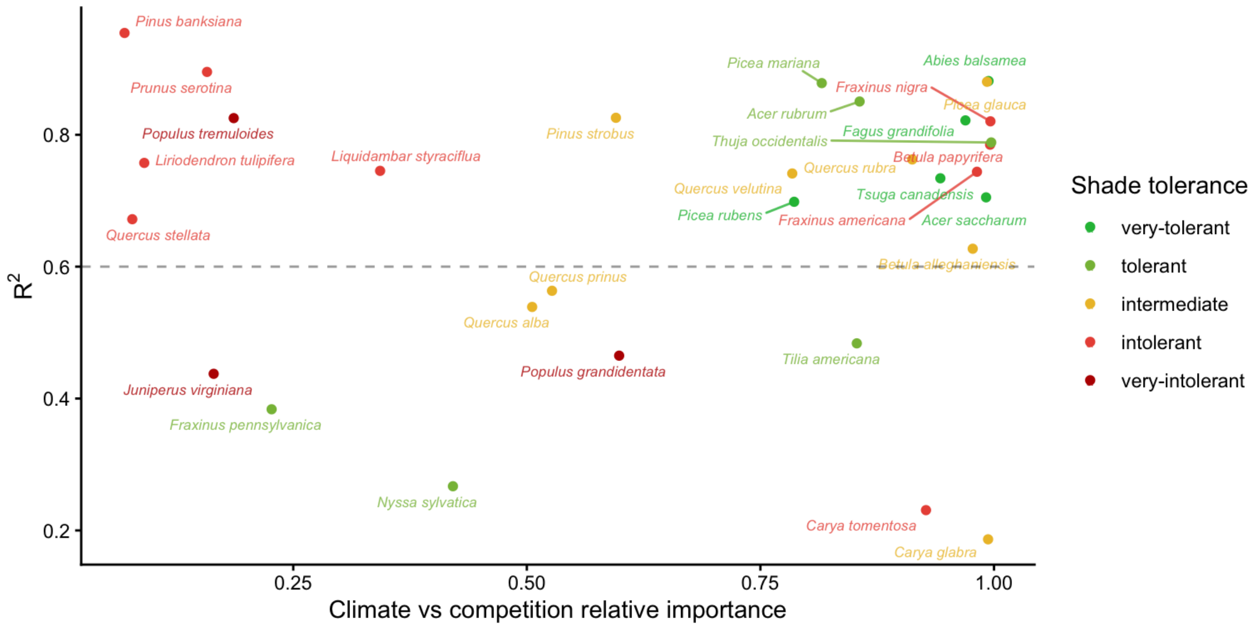

Figure 24: Distribution of relative importance between climate and competition covariates according to Random Forest, the respective $R^2$. The more species are to the right of the panel, the more climate is important relative to competition. Color represents the level of shade tolerance (Burns et al. 1990)

variation in $\lambda$, previously attributed to the growth model, shifted to the recruitment model. Also, the importance attributed to the survival model for certain species at the center and cold temperature conditions shifted toward the recruitment model. Although we observed this shift, the overall patterns remained similar to those discussed earlier. The only exception was the distribution of relative importance between climate and competition (Figure 24), where many species had an increase in the importance of competition relative to climate. These observed differences primarily arise from the high sensitivity of $\lambda$ to the $\phi$ parameter.

### References

Antoniadis, A., S. Lambert-Lacroix, and J.-M. Poggi. 2021. Random forests for global sensitivity analysis: A selective review. Reliability Engineering & System Safety 206:107312.

Breiman, L. 2001. Random forests. Machine learning 45:5–32.

Burns, R. M., B. H. Honkala, and Others. 1990. Silvics of North America: 1. Conifers; 2. Hardwoods Agriculture Handbook 654. US Department of Agriculture, Forest Service, Washington, DC.

Caswell, H. 1978. A general formula for the sensitivity of population growth rate to changes in life history parameters. Theoretical population biology 14:215–230.

Díaz, S., J. Kattge, J. H. C. Cornelissen, I. J. Wright, S. Lavorel, S. Dray, B. Reu, M. Kleyer, C. Wirth, I. C. Prentice, and Others. 2022. The global spectrum of plant form and function: enhanced species-level trait dataset. Scientific Data 9:755.

Gabry, J., R. Češnovar, and A. Johnson. 2023. cmdstanr: R Interface to 'CmdStan'.

Lewontin, R. C., and D. Cohen. 1969. On population growth in a randomly varying environment. Proceedings of the National Academy of sciences 62:1056–1060.

Saltelli, A., K. Aleksankina, W. Becker, P. Fennell, F. Ferretti, N. Holst, S. Li, and Q. Wu. 2019. Why so many published sensitivity analyses are false: A systematic review of sensitivity analysis practices. Environmental modelling & software 114:29–39.

Team, S. D., and Others. 2022. Stan modeling language users guide and reference manual, version 2.30.1. Stan Development Team.

Vehtari, A., A. Gelman, and J. Gabry. 2017. Practical Bayesian model evaluation using leave-one-out cross-validation and WAIC. Statistics and computing 27:1413–1432.

Wright, M. N., and A. Ziegler. 2017. {ranger}: A Fast Implementation of Random Forests for High Dimensional Data in {C++} and {R}. Journal of Statistical Software 77:1–17.
